# Perceiving Depth from Texture and Disparity Cues: Evidence for a Non-Probabilistic Account of Cue Integration

**DOI:** 10.1101/2022.10.20.513044

**Authors:** Jovan T. Kemp, Evan Cesanek, Fulvio Domini

## Abstract

The fundamental question of how the brain derives 3D information from the inherently ambiguous visual input has been approached during the last two decades with probabilistic theories of 3D perception. Probabilistic models, such as the Maximum Likelihood Estimation (MLE) model, derive from multiple independent depth cues the most probable 3D interpretations. These estimates are then combined by weighing them according to their uncertainty to obtain the most accurate and least noisy estimate. In three experiments we tested an alternative theory of cue integration termed the Intrinsic Constraint (IC) theory. This theory postulates that the visual system does not derive the most probable interpretation of the visual input, but the most stable interpretation amid variations in viewing conditions. This goal is achieved with the Vector Sum model, that represents individual cue estimates as components of a multidimensional vector whose norm determines the combined output. In contrast with the MLE model, individual cue estimates are not accurate, but linearly related to distal 3D properties through a deterministic mapping. In Experiment 1, we measured the cue-specific biases that arise when viewing single-cue stimuli of various simulated depths and show that the Vector Sum model accurately predicts an increase in perceived depth when the same cues are presented together in a combined-cue stimulus. In Experiment 2, we show how Just Noticeable Differences (JNDs) are accounted for by the IC theory and demonstrate that the Vector Sum model predicts the classic finding of smaller JNDs for combined-cue versus single-cue stimuli. Most importantly, this prediction is made through a radical re-interpretation of the JND, a hallmark measure of stimulus discriminability previously thought to estimate perceptual uncertainty. In Experiment 3, we show that biases found in cue-integration experiments cannot be attributed to flatness cues, as assumed by the MLE model. Instead, we show that flatness cues produce no measurable difference in perceived depth for monocular (3A) or binocular viewing (3B), as predicted by the Vector Sum model.

## Introduction

A fundamental aspect of human visual perception is its ability to interpret three-dimensional space from patterns of light. We may be able to ignore color when judging brightness or divert our attention from specific objects with eye movements, but we cannot possibly suppress our experience of a three-dimensional environment. The problem of how the visual system constructs a 3D interpretation from the two-dimensional manifold of light intensity at the retina has been approached during the last three decades through a probabilistic inference theory of 3D vision (Landy et al., 2011; Landy et al., 1995). The intuitive appeal of this theory has led to a large number of empirical studies aimed at evaluating its predictions (Adams et al., 2004; Adams & Mamassian, 2004; Ernst & Banks, 2002; Chen, & Saunders, 2019; Jacobs, 1999; Jacobs, 2002; Knill, 1998a; Knill, 2007; Knill & Saunders, 2003; Mamassian & Landy, 1998; Hillis et al., 2002; Hillis et al., 2004; Saunders & Chen; 2015; Schrater & Kersten, 2000; Saunders & Knill, 2001; Welchman et al., 2008; Young et al., 1993). Though this approach successfully accounts for a wide range of findings, it is unable to predict many fundamental real-world phenomena, such as systematic biases in 3D judgments (Bozzacchi & Domini, 2015; Bozzacchi et al., 2016; Campagnoli et al., 2017; Caudek & Domini, 1998; Domini & Braunstein, 1998; Domini & Caudek, 1999; Domini & Caudek, 2003; Domini et al., 1998; Egan & Todd, 2015; Fantoni et al., 2010; Kopiske et al., 2019; Liu & Todd, 2004; Norman et al., 2004; Norman et al., 1996; Norman et al., 1995; Perotti et al., 1998; Phillips & Todd, 1996; Tittle et al., 1995; Todd, 2004; Todd & Bressan, 1990; Todd et al., 1998; Todd & Norman, 2003; Todd et al., 2014; Todd & Thaler, 2010; Todd et al., 2005; Todd et al, 2007; Todd et al., 1995; Volcic et al, 2013), internal inconsistencies among judgments at different scales (Lappin & Craft, 2000; Loomis et al., 1996; Loomis et al., 2002), the paradox of pictorial depth and pictorial duality (Haber, 1980; Koenderink, 1998; Koenderink et al., 2001; Vishwanath, 2011; 2013; 2014; 2020), and differences in phenomenology of 3D vision (Koenderink et al., 2015; Koenderink et al., 2018; Vishwanath, 2013). In this paper, we test a new theoretical framework based on an entirely different set of assumptions that can more parsimoniously account for the full range of observations in 3D perception.

There are two main assumptions that have guided recent research in 3D vision: (1) Independent modules derive noisy estimates that are on average veridical (i.e. unbiased) (Clark & Yuille, 1990; Landy et al., 2011) and (2) visual mechanisms also estimate the magnitude of sensory noise, such that the outputs of individual modules represent probability distributions. Representing probability distributions enables the statistically optimal combination of independent estimates, as proposed by Bayesian integration frameworks (*e*.*g*., Landy et al., 2011). Although there are more general implementations of Bayesian combination, in this paper we focus on the linear Maximum Likelihood Estimation (MLE) model (Ernst & Bülthoff, 2004), following similar past studies that have assumed a negligible influence of priors when viewing objects defined by binocular disparity, texture, or both (Chen & Saunders, 2020; Hillis et al., 2004; Johnston et al., 1993; Knill & Saunders, 2003).

The predictions of the linear MLE model for the integration of texture and disparity information can be summarized by two equations. First, if 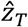and 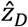are the depth estimates from the texture and disparity modules and *σ*_*T*_ and *σ*_*D*_ are the standard deviations of the noise of these estimates, then the combined estimate 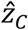 is a weighted average with weights proportional to the reliabilities of the estimates:

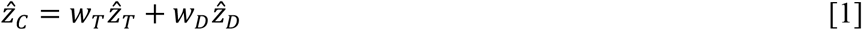

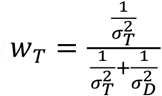 and 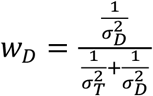. Second, the variance of the combined estimate is smaller than that of either single-cue estimate, as predicted by the following relationship:

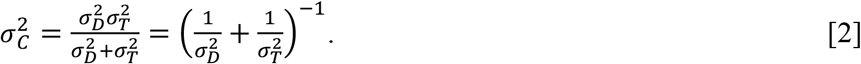

While applying the MLE model to explain perceptual processing may appear straightforward, some of its core assumptions seem not to be satisfied by human perceptual systems. First, many experiments have shown that texture, motion, and binocular disparity cues generally fail to produce accurate percepts, contrary to the veridicality assumption (Bozzacchi & Domini, 2015; Bozzacchi et al., 2016; Campagnoli et al., 2017; Caudek & Domini, 1998; Domini & Braunstein, 1998; Domini & Caudek, 1999; Domini & Caudek, 2003; Domini et al., 1998; Egan & Todd, 2015; Fantoni et al., 2010; Kopiske et al., 2019; Liu & Todd, 2004; Norman et al., 2004; Norman et al., 1996; Norman et al., 1995; Perotti et al., 1998; Phillips & Todd, 1996; Tittle et al., 1995; Todd, 2004; Todd & Bressan, 1990; Todd et al., 1998; Todd & Norman, 2003; Todd et al., 2014; Todd et al., 1995; Todd & Thaler, 2010; Todd et al., 2005; Todd et al., 2007; Volcic et al., 2013). Second, it has been shown that when perception is measured with techniques other than depth discrimination (*e*.*g*. by setting an independent 2D probe), the measured variability in perceived depth does not predict the relative weighting of depth cues (Todd et al., 2010), contrary to the assumption that cue estimates are represented as probability distributions. These considerations suggest that the widespread application of the MLE model to human 3D perception may be inappropriate, and that cue-combination experiments need to be reinterpreted with an alternative explanation.

Here we aim to develop a theoretical framework of 3D cue combination that does not require any of the controversial assumptions of the mainstream MLE account described above. Instead, this framework assumes: (1) A derivation of estimates of 3D properties that are generally biased but under some viewing conditions may be veridical and (2) are deterministic rather than probabilistic estimates of 3D properties from single and multiple signals. The combination rule for multiple signals is therefore not dependent on knowledge of variance within a cue estimate. Instead, this process is optimized to achieve perceptual stability in face of the natural variation of viewing conditions and material composition of external surfaces. In the next section, we provide a formal specification of this framework that makes specific quantitative predictions. We then test these predictions in three experiments. Notably, we find that the model accurately predicts the reduction in discrimination threshold that occurs when additional depth cues are added to a stimulus. This finding has been interpreted as a critical piece of evidence for the MLE model, but here we show that it is entirely consistent with our novel framework. Moreover, our model predicts several novel results that cannot be predicted by previous theories of cue integration.

### Intrinsic Constraint Theory of multi-cue processing

The computational model we propose is termed the Intrinsic Constraint (IC) theory, in reference to the original model from which it was developed (Domini & Caudek, 2009; Domini et al., 2006). Importantly, however, it is based on entirely different assumptions than the earlier IC theory.

First, we postulate that separate visual modules independently process distinct image regularities. We assume these modules are tuned to approximate a linear mapping between the distal 3D property *z* and the internal 3D estimate 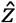. The slope of this linear function depends on the strength of the visual information. For instance, from the image in Figure 1, a texture module extracts the systematic change in shape and spatial frequency of texture elements resulting in an estimate 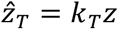. The IC theory defines visual modules as independent insofar as the slopes *k* of the transfer functions vary independently. Critically, notice that in direct contrast to the MLE model, there is no assumption that the transfer function is veridical, nor is there any explicit representation of the associated sensory noise. These simplifications make the IC theory far more parsimonious than the MLE model.

**Figure 1:**
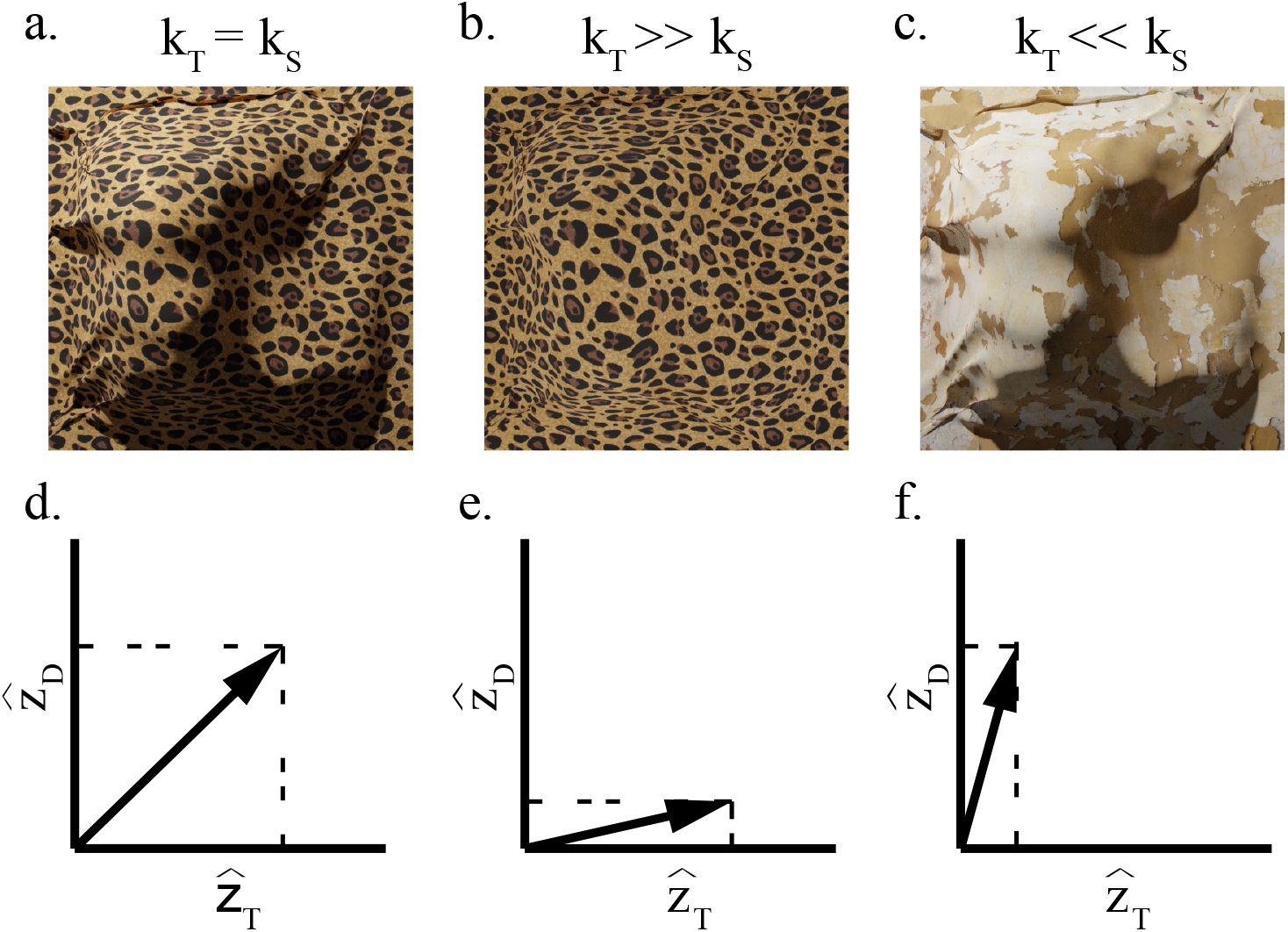
(**a-c**) A series of bumpy surfaces with varying levels of texture and shading gradients. The cue strength, k_*i*_, depends on the “quality” of the gradients, with more detectible gradients producing larger cue strength. Textures with identifiable, regular elements and shading gradients with continuously varying luminance intensities produce large cue strengths that in this example are assumed to be identical (**a**). Textures with ambiguous elements (**c**) and shading with small luminance differences (**b**) produce smaller cue strengths. (**d-e**) Each cue is analyzed by an independent function that produces a depth signal, which is linearly proportional to the distal depth. These signals exist in a multidimensional space where the vector length of the signals depend on the depth of the surface and the cue strength of the cue. It is this vector length which drives perceived depth magnitude. If both cues form strong gradients (**d**), then the vector length and subsequent perceived depth will be large. Weaker gradients for either cue will reduce this vector length and the perceived depth (**e, f**).

To illustrate the independence of the slopes of the transfer functions, consider the bumpy surface depicted in Figure 1a. Now imagine how the various image signals indicating the 3D relief of this surface would be affected by different viewing conditions. For instance, overcast weather would wash out the shading gradient while leaving the texture pattern unmodified, resulting in the image shown in Figure 1b. On the other hand, the same surface can be covered with irregular texture rather than the highly regular cheetah pattern while the lighting condition remains the same. The shading pattern would be unmodified but there would be no clear gradient of texture elements, as depicted in Figure 1c. In these examples, independent confounding variables are associated with the material composition of the surface and the sources of illumination. In a similar fashion, different confounding variables will affect other image signals, such as the speed of the observer in motion parallax (Fantoni et al., 2012) or the fixation distance between the observer and the object in binocular disparities (Johnston, 1991). Thus, in general, the slopes of the transfer functions for modules devoted to processing different image regularities will vary independently across stimuli and viewing contexts. Note that within this framework, a type of visual information traditionally defined as a single, monolithic cue may in fact be better understood as multiple cues, so long as they are affected by independent confounding variables. Indeed, by this definition there are several distinct types of texture cues that are traditionally treated as a single cue (Chen & Saunders, 2020; Todd & Thaler, 2010; Todd et al., 2007).

Since the slopes of the transfer functions, *k* are determined by the strengths of individual cues, we will refer to these parameters as *cue strengths*. As a consequence of independent cue strengths, we can represent the totality of the cue estimates derived from a given stimulus as a multidimensional vector. This is illustrated in Figures 1d-f, which correspond to three stimuli composed of texture and shading information (Figs. 1a-c). Figure 1d depicts a stimulus for which texture and shading (arbitrarily) have the same strength (*k*_*T*_ = *k*_*S*_). In Figure 1e, the strength of texture is much greater than the strength of shading, as the shading has been removed (*k*_*T*_ ≫*k*_*S*_). In Figure 1f, the strength of shading is much greater than the strength of texture, as the texture is highly irregular (*k*_*T*_ ≪ *k*_*S*_). Since both 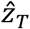 and 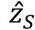 are proportional to the 3D property z, the length of the combined vector 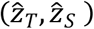 is also proportional to z. However, since the combined vector length depends on the individual cue strengths, it will therefore fluctuate with the confounding variables. *Critically, the central claim of our theory is that the goal of the visual system is to maximize sensitivity to underlying 3D information while minimizing sensitivity to confounding variables*.

In Appendix 1, we show that the combined estimate, calculated as a vector-sum of single-cue signals scaled by parameters representing the variability of independent confounding variables, achieves this goal. For the general case of multiple image signals this leads to the Vector Sum Equation:

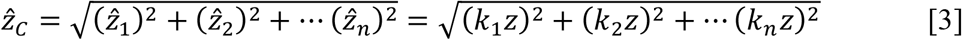

There are several nuances to the IC theory that should be noted. First, the value *z* does not necessarily imply a single depth value or depth map, but more generally can also signify a slant or a curvature map. Second, the cue strengths *k*_*i*_ are not free parameters, they are the empirically identified values for the single-module transfer function. Third, since the model is additive, it may be misunderstood as producing systematic overestimations as more cues are added to a stimulus. In fact, it is quite the contrary. We speculate that removing cues brings the model outside of its optimal operating conditions, which results in underestimation of depth from reduced- or single-cue stimuli. Note that we will refer to the phenomenon of observing an increase in depth with the addition of cues as the *Vector Sum* model.

Another important point to highlight is about the source of variability of 3D estimates that is considered relevant from the IC theory perspective. Previous models assume that depending on the “quality” (i.e., reliability) of 3D information specifying a given stimulus, 3D estimates will fluctuate from one view of the stimulus to the next. For instance, multiple views of stimuli carrying the same regular texture pattern of Figure 1b. will produce a much smaller variation of 3D judgements than repeated viewing of the plaster texture of Figure 1c. In contrast, the IC theory only predicts negligible fluctuations, due to unavoidable neural noise and slight variations in the texture patterns from one view to the next. The relevant variability of 3D estimates affecting repeated viewing of the *same distal structure* is instead due to a change of the confounding variables (e.g. the material composition of the object, resulting in a change of the strength of the texture pattern). What is fundamental to this theory is that the Vector Sum combination rule is blind to the strength of each individual cue. Therefore, it does not, as MLE models, dynamically weigh the output of single-cue modules according to their individual “quality”.

The main goal of this study is to test the efficacy of the Vector Sum model in predicting several documented properties of depth perception while reinterpreting the mechanisms which bring about cue processing and combination. Experiment 1 examines the inaccuracy of single-cue estimates and the systematic biases that can be expected when cues are combined. We show that these biases can be predicted without free parameters through the Vector Sum Model. Experiment 2 replicates the previous finding that discrimination thresholds decrease for combined-cue stimuli relative to single-cues. We discuss why the Vector Sum model and the MLE model make similar predictions regarding discrimination thresholds, but for very different reasons (*i*.*e*., reasons related to the properties of linear cue strengths versus the properties of probability distributions). Experiment 3 provides evidence that cues-to-flatness are unlikely to allow the MLE model to account for the biases that the Vector Sum model successfully predicts.

## Experiment 1

A first test of the Intrinsic Constraint theory is to verify that the combination of multiple cues leads to depth estimates in alignment with the Vector Sum Model. According to Equation 3, the perceived depth of a combined-cue stimulus is predicted to be larger than the perceived depth of single-cue stimuli. For the specific case of texture and binocular disparities Equation 3 can be reduced to the following equation:

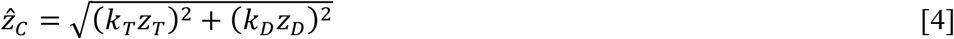

In single-cue conditions only one cue is present, which means that the cue strength of all absent cues is zero. We studied the perceived depth of a sinusoidally corrugated surface by manipulating the amplitude of the sinusoid. In the disparity-only condition, the surface was specified by a random-dot stereogram (RDS) which did not provide any discernible texture information (i.e., *k*_*T*_ = 0). In the texture-only condition, a compelling texture gradient specified the depth profile of the surface while binocular disparities were set to zero (*z*_*D*_ = 0; equivalent to *k*_*D*_ = 0 in the Vector Sum model). The choice of rendering the texture-only stimulus binocularly was made for the practical reason of keeping the vergence signal constant in all viewing conditions. In the combined-cue condition both texture and disparity information were present in the stimulus.

### Experiment 1: Methods

#### Participants

Eleven participants (3 being the authors) were drawn from the Brown University community and participants completed Experiment 1. Participants either received $12/hour or course credit as compensation. Participants provided informed consent prior to testing. The procedure reported was approved by the Brown University Institutional Review Board.

#### Apparatus

Experiments were completed on a Dell Precision T7500 powered by a nVidia Quadro 4000 graphics card. Stimuli were simulated on a Sony Triniton GDM-f520 CRT monitor with a resolution of 1280×1024 at a refresh rate of 85hz. The display was projected onto a half-silvered mirror that was slanted 45 deg about the vertical axis in front of the participant with respect to the fronto-parallel plane. The monitor was repositioned to different viewing distances via a Velmex linear actuator (Velmex, Inc., Bloomfield, NY).

Binocular disparity was provided using NVIDIA 3D Vision® 2 wireless glasses (NVIDIA, Santa Clara, CA) which were synchronized to the refresh rate of the monitor to provide unique images to each eye. The interocular distance (*IOD*) of every participant was measured using a digital pupillometer (Reichert Inc., Depew, NY). Participants viewed the stimuli while positioned on a chinrest.

#### Stimuli

The target stimuli were three-dimensional corrugated surfaces whose depth profile followed a sinusoidal modulation along the vertical axis. An example stimulus and probe presented to participants is shown in Figure 2a. The corrugated surface was seen through a square frame subtending approximately 8° of visual angle to eliminate contour information. The wave period was 4.50° of visual angle.

**Figure 2:**
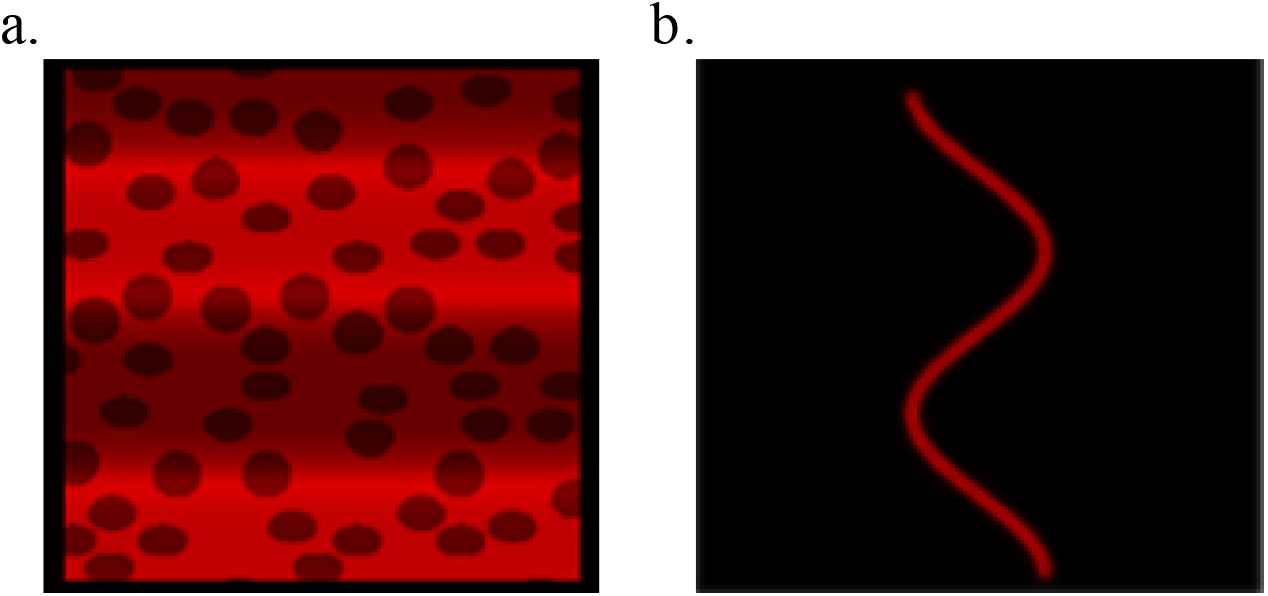
An example of a monocular view of the sinusoidal stimulus (**a**) and the 2D probe (**b**). The surface of the stimulus is defined by shading and texture information. This combination of depth cues is referred to as “texture cue” for simplicity. In the combined-cue condition the 3D structure was also specified by binocular disparities. Participants adjusted the amplitude of the 2D probe to report perceived depth. In this example, the probe is set to the correct amplitude which matches the simulated depth profile of the surface.

Participants made depth judgments by adjusting a two-dimensional sinusoidal probe whose horizontal amplitude varied along the vertical axis. The wave period of the probe also subtended 4.50° of visual angle. An example of the probe with its amplitude set to the correct magnitude is shown below the corrugated surface in Figure 2b. The phase of the 3D surface was randomly varied on each trial to eliminate depth adaptation. However, the phase of the probe line remained constant throughout all sessions.

Participants judged the depth of three types of 3D information: texture-only, disparity-only, and combined-cue stimuli. Texture-only sine waves were constructed by volumetric texturing. This process first involved randomly placing the centers of spheres with radii subtending visual angle of 0.55° onto the simulated 3D corrugated surface. Any portion of the wave that intersected a sphere was darkened relative to the surrounding red surface. This produced a compelling texture gradient on the image projection. To eliminate depth order ambiguity, shading information was produced by placing a single directional light source from above oriented at a 45° with respect to the fronto-parallel plane. We refer to this as texture cue for simplicity. To keep a steady fixation at the center of the display as in the disparity-only and combined-cue conditions, texture-only stimuli were also seen binocularly.

Disparity-only surfaces were constructed with a random dot stereogram consisting of 400 dots each with a visual angle of 0.1°. The dots were uniformly distributed on the image plane with the constraint that they did not overlap. Because part of the surface was occluded by a frame, there were on average 320 visible dots. Two views were rendered by placing the rendering cameras at the estimated locations of the observer’s nodal points, which were determined after measuring for each observer the interocular distance (IOD). NVIDIA 3D Vision® 2 wireless stereo-glasses were used to separate the projection of the left and right images for the appropriate eye. Combined-cue stimuli were obtained by rendering stereoscopically the polka-dot textured surfaces.

#### Procedure

Participants completed two blocks within a single session. Each block had a constant fixation distance of either 40 or 80 cm. Within each block, participants viewed sinusoidal surfaces with four different peak-to-trough depths (2.5, 5, 10, or 15 mm) defined by one of three cue types (disparity-only, texture-only, and combined-cue), with 7 repetitions for each combination of depth and cue type. Thus, each block involved 84 judgments and lasted approximately 20 minutes. At the onset of each trial, a fixation cross was displayed for 700 ms, followed by the presentation of the surface stimulus, as well as a 2D sine wave probe icon at the bottom of the display (Fig. 2b). Participants adjusted the amplitude of the icon until the peak-to-trough length matched their perceived depth of the target stimulus. During the adjustment they were free to move their eyes back and forth between the 3D surface and the 2D icon. Once they were satisfied with their setting, they submitted their judgment with a button press, which also initiated the next trial. Before the experimental session, participants completed a small number of practice trials with stimuli of random depths. No feedback about response accuracy was provided at any point.

### Experiment 1: Results and Discussion

Qualitatively, the Vector Sum model predicts that the combined-cue stimulus should be perceived deeper than the single-cue stimuli. Figure 3 shows the average probe settings across cues (denoted by line color) and fixation distances (denoted by separate panels). A repeated-measures ANOVA found a main effect of simulated depth (*F*(1,10) = 272.67, *p =* 1.4e-8 ; Generalized *η*^2^ = 0.89) and cue type (*F*(2,10) = 48.40, *p* = 2.2e-8; Generalized *η*^2^ = 0.43). For both fixation distances, the perceived depth of combined-cue stimuli (purple diamonds) was consistently greater than the perceived depth of single-cue stimuli (red squares, blue circles). A Bonferroni-corrected post-hoc analysis confirmed that perceived depth in the combined-cue condition was larger than the perceived depth in both the disparity-only condition (*T*(10) = 4.42, *p* = 0.0039) and the texture-only condition (*T*(10) = 10.59, *p* = 2.8e-6). Additionally, texture-only stimuli were in general perceived as shallower than disparity-only stimuli, demonstrating cue-specific biases (*T*(10) = -5.02, *p* = 0.0016).

**Figure 3.**
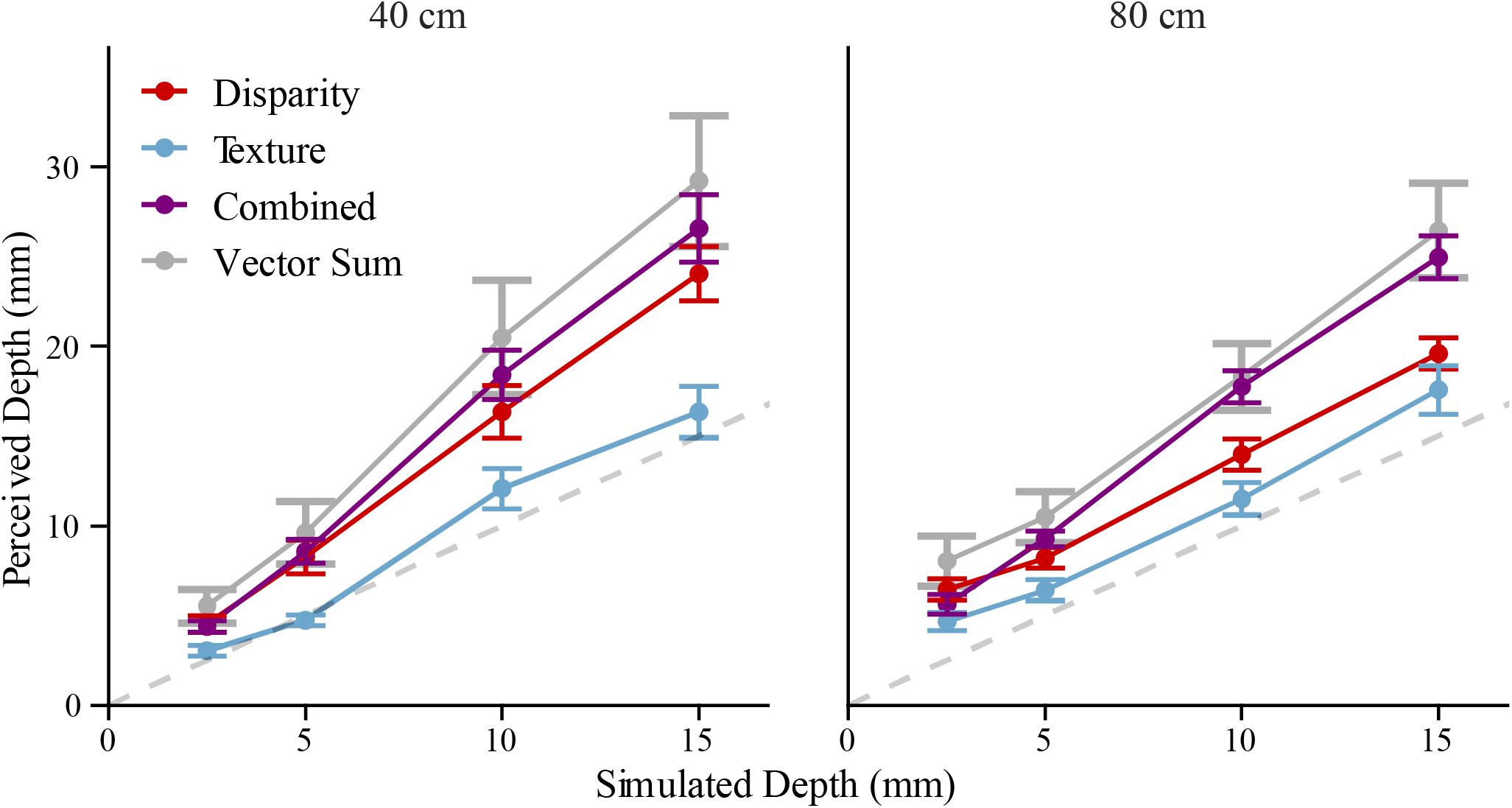
Average depth judgments as function of simulated depth for viewing distances of 40 (left panel) and 80 cm (right panel) with error bars showing the standard error of the mean. The horizontal axis labeled “Simulated Depth” represents the peak to trough depth of the corrugated 3D surface while the vertical axis labeled “Perceived Depth” represents the set amplitude of the 2D probe. The different cue conditions are denoted by the shape and color of the data points: purple squares for the combined-cue condition, red diamonds for the disparity condition, and blue triangles for the texture condition. The Vector Sum prediction is denoted by the grey line with 95% confidence intervals at each point. The dashed grey line is the unity line denoting veridical perception.

All interactions were significant. The interaction between cue type and fixation distance (*F*(2,20) = 3.76, *p* = 0.041; Generalized *η*^2^ = 0.03), between simulated depth and fixation distance (*F*(1,10) = 7.19, *p* = 0.023; Generalized *η*^2^ = 0.07), and between all three factors (*F*(2,20) = 5.11, *p* = 0.016; Generalized *η*^2^ = 0.031) reflects the dependence of cue strength on how the fixation distance influences the quality of the cue. This was expected particularly for the disparity-only cue where a lack of depth constancy across distances is a well-documented phenomenon (Johnston, 1991). The interaction between simulated depth and cue type (*F*(2,20) = 45.42, *p* = 3.7e-8; Generalized *η*^2^ = 0.20) further supports the existence of cue-specific biases due to differing cue strengths between cue types.

The Vector Sum model predicts that the perceived depth of the combined cue should be the square root of the sum of squares of the perceived depth of the single-cues (eqs. 3 and 4). Figure 3 plots the average predictions of the Vector Sum model across participants with 95% confidence intervals (gray). Given that we assume a (ideally) linear mapping, the model can directly predict the cue strength of the combined-cue from those of the single-cues through Equation 4. The prediction is simplified to the following since the simulated depth rendered for each cue is the same (*i*.*e*., there are no cue conflicts):

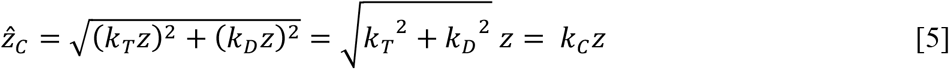

Since the slopes of the functions relating perceived to distal depth are proxies for the cue strengths, Equation 5 predicts the slope of the combined-cue estimate 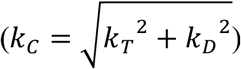 from the slopes of the single-cue estimates (*k*_*T*_ and *k*_*D*_) without any free parameters. Figure 4 shows the predicted slopes plotted against the measured slopes for each participant. The correlation coefficient *r* was found to be 0.79 while a linear fit with an intercept of zero found a slope of 0.96 (*SE =* 0.034) showing a close match to the unity line.

**Figure 4:**
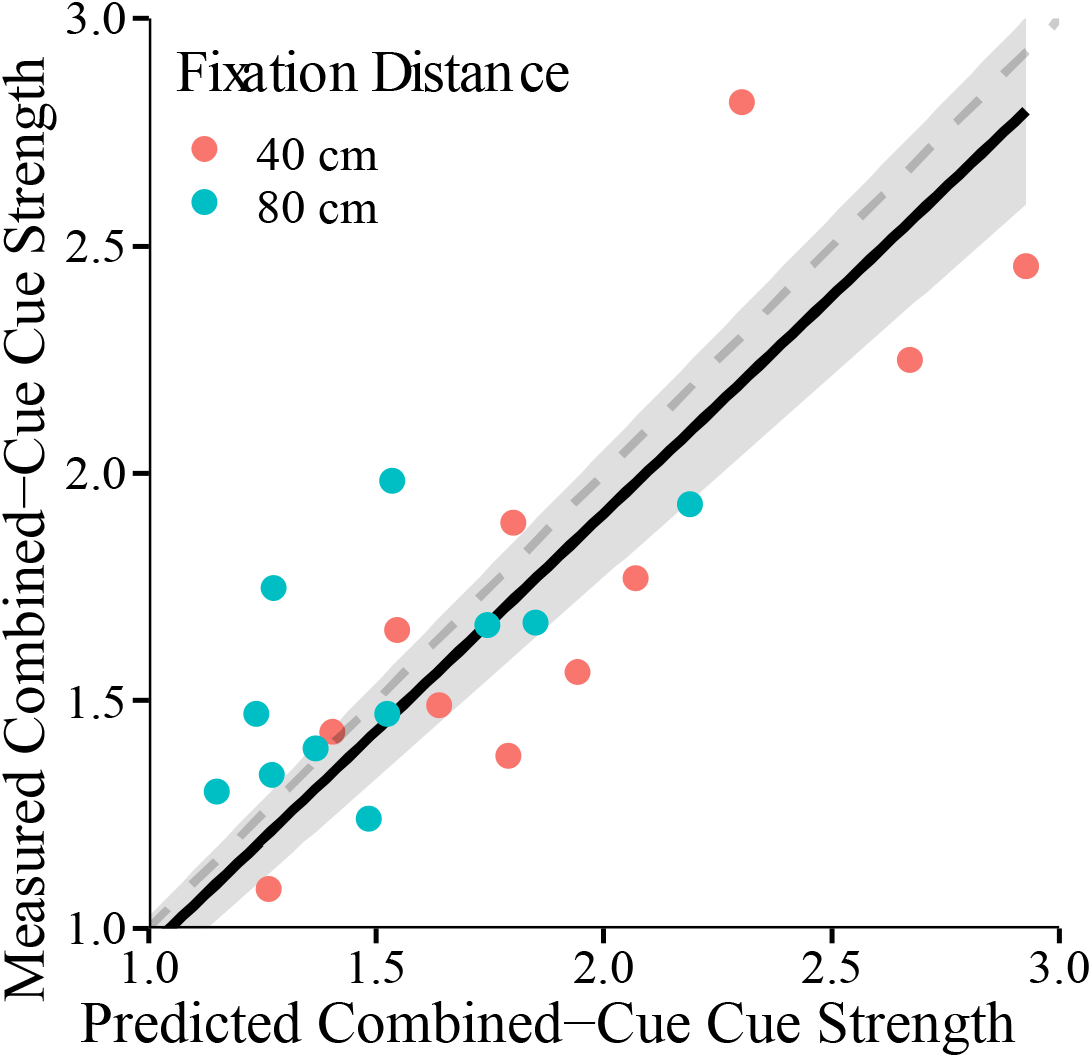
Observed combined-cue strength vs. predicted combined-cue strength. The predicted combined-cue strength is computed with the Vector Sum model without free parameters directly from the single-cue strengths. The single-cue and combined-cue strengths were determined by the slopes of linear fits. Each data point represents a subject either in the 40 cm (red) or 80 cm (blue) fixation distance condition. The dashed grey line represents accurate prediction. The gray area denotes the 95% confidence interval of the linear fit (black line) of the observed vs. the predicted cue-strength.

Overall, these results demonstrate that the Vector Sum Model produces highly accurate predictions of the relationship between simulated and perceived depth in single- and combined-cue conditions, with no free parameters. In contrast, the results clearly contradict the MLE model prediction that the combined-cue perceived depth will fall between the single-cue perceived depths. Although the MLE model predictions may be amended by introducing cues-to-flatness, we will provide evidence in Experiment 3 rejecting the cues-to-flatness explanation. Additionally, single-cue and combined-cue depths were consistently overestimated in five of six stimulus conditions, contradicting the veridicality assumption of the MLE model.

If previous findings from ostensibly similar tasks have supported the MLE model (Hillis et al., 2004; Knill & Saunders, 2003; Lovell et al., 2012) then why does the MLE model fail in predicting these results? A critical difference is that observers in this task provided absolute judgments of depth using a probe figure, whereas in earlier studies observers made relative judgments by comparing or matching two 3D shapes.

While relative judgment tasks are often useful, they cannot reveal systematic biases in depth perception. For example, Hillis et al. (2004) asked participants to match the perceived slant of an adjustable cue-consistent surface with the slant of a fixed cue-conflict surface (*i*.*e*., the simulated slants from texture and from disparity were either matched or mismatched). The cue-consistent slant that yielded a match was predicted through the MLE model (Equation 1). However, there is no guarantee that either surface was perceived veridically. Nevertheless, it is notable that discrimination thresholds measured on single-cue stimuli were indeed good predictors of the weights estimated in the slant matching task. The IC theory, however, provides a radically different interpretation of discrimination thresholds. When this new interpretation is adopted it can be shown that an approximation of the Vector Sum model makes identical predictions of the results of Hillis et al. (2004) to those of the MLE model (Appendix 2).

#### Cue Uncertainty and Judgment Variance

An important prediction of the MLE model is that the variance of the combined-cue estimate should be smaller than the variances of the single-cue estimates (eq. 2). Test of this MLE prediction is usually conducted by measuring discrimination thresholds of single-cue and combined-cue stimuli. However, noise coming from depth estimation should also surface in the standard deviation of probe adjustments. We should therefore expect that the standard deviation of probe adjustments in the combined-cue condition should be smaller than that measured in the single-cue conditions. Alternatively, the Vector Sum model assumes that depth estimates are basically deterministic, only affected by negligible neural noise. According to this theory variability in perceptual judgements is all due to late-stage, task related processes independent of the stimulus itself. We therefore should expect that there is no difference between the cue types for the response variance. Given the different predictions of the two models, we tested whether there was a difference in the SD between the cues. Figure 5 shows the standard deviation of the probe-adjustment task as function of simulated depth in all experimental conditions. In this figure the prediction of the MLE model for the standard deviation of the combined-cue adjustments is shown in gray.

**Figure 5:**
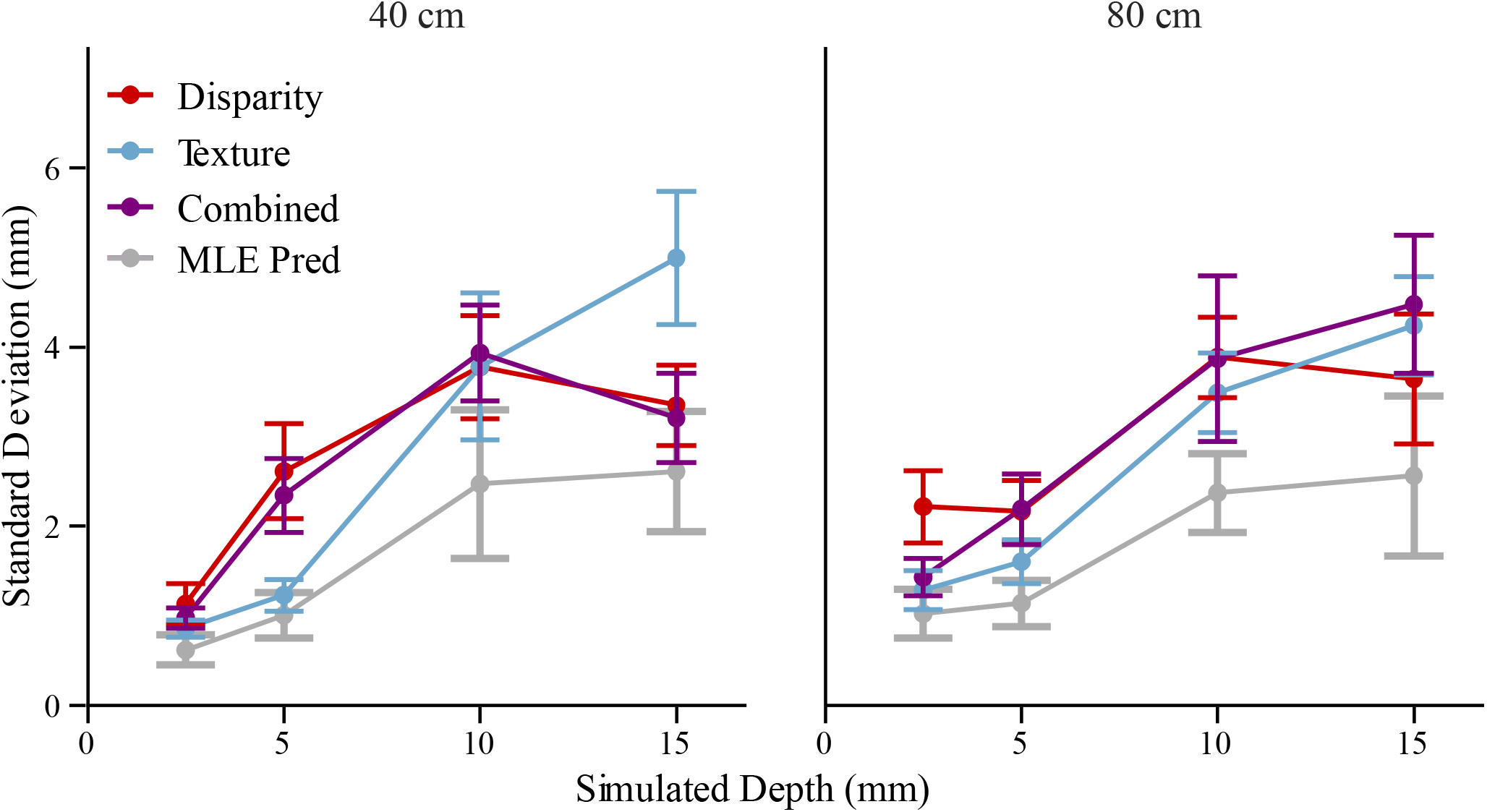
The average standard deviations of the probe adjustment task in Experiment 1 with error bars averaged across subjects. The MLE predictions together with the 95% confidence intervals are shown in gray. The do not align with the MLE predictions since the SDs observed in the combined-cue condition are not smaller the *SD*s observed in the single-cue conditions.

A repeated measures ANOVA indicated one main effect of simulated depth (*F*(1,10) = 54.75, *p* = 2.3e-5; Generalized *η*^2^ = 0.57). This follows the classic effect of Weber’s law where the response variance is proportional to the magnitude of the stimulus, in this case the surface depth. There was also an interaction between the cue type and simulated depth (*F*(2,20) = 7.31, *p* = 0.0041; Generalized *η*^2^ = 0.09). However, there was no main effect of cue type (*F*(2,20) = 0.45, *p* = 0.65; Generalized *η*^2^ = 0.0053). This can be easily observed in Figure 5 where the combined-cue standard deviation (purple) is not smaller than the single-cue standard deviations, as predicted by the MLE model (gray). Instead, these results support the prediction of the Vector Sum model that noise observed in perceptual judgements is stimulus independent. Because the Vector Sum predicts a null effect of cue type, we conducted a Bayes factor analysis using the *BayesFactor* package in R (Morey & Rouder, 2021). A Bayes factor of 0.055 indicated strong evidence for a model including fixed effects of simulated depth and fixation distance, compared to a model including the same fixed effects with the inclusion of cue type. Both models included a random effect for participants.

These results are particularly intriguing since they seem to be inconsistent with findings obtained in experiments where discrimination thresholds are used to test the predictions of the MLE model. Indeed, results from discrimination threshold experiments suggest that the variance of combined-cue stimuli is smaller than the variance of single-cue stimuli by an amount predicted by Equation 2. This quantitative prediction, however, is also compatible with the prediction of the Vector Sum model once discrimination thresholds are interpreted in a radically different way.

## Experiment 2

The central hypothesis of the MLE framework is that cue combination leads to an increase in the reliability of the depth estimate. In many previous investigations, the reliability of a depth estimate has been assumed to be directly reflected by the just-noticeable difference (JND) in a two-interval forced choice (2IFC) task. The JND is the difference in distal depth that leads to 84% accuracy in identifying the deeper stimulus. Under the MLE model, this is interpreted as the standard deviation of the noise in the estimation process. Figure 6a depicts how in typical MLE models JNDs arise from a noise-free decision process that compares two noisy estimates. For example, the JNDs are larger for a disparity stimulus at near viewing distances than at far viewing distances due to less estimation noise.. Studies using this approach have repeatedly demonstrated that single-cue and combined-cue JNDs adhere to the relationship predicted by the MLE model (eq. 2; Ernst & Banks, 2002; Hillis et al., 2004; Knill & Saunders, 2003).

**Figure 6:**
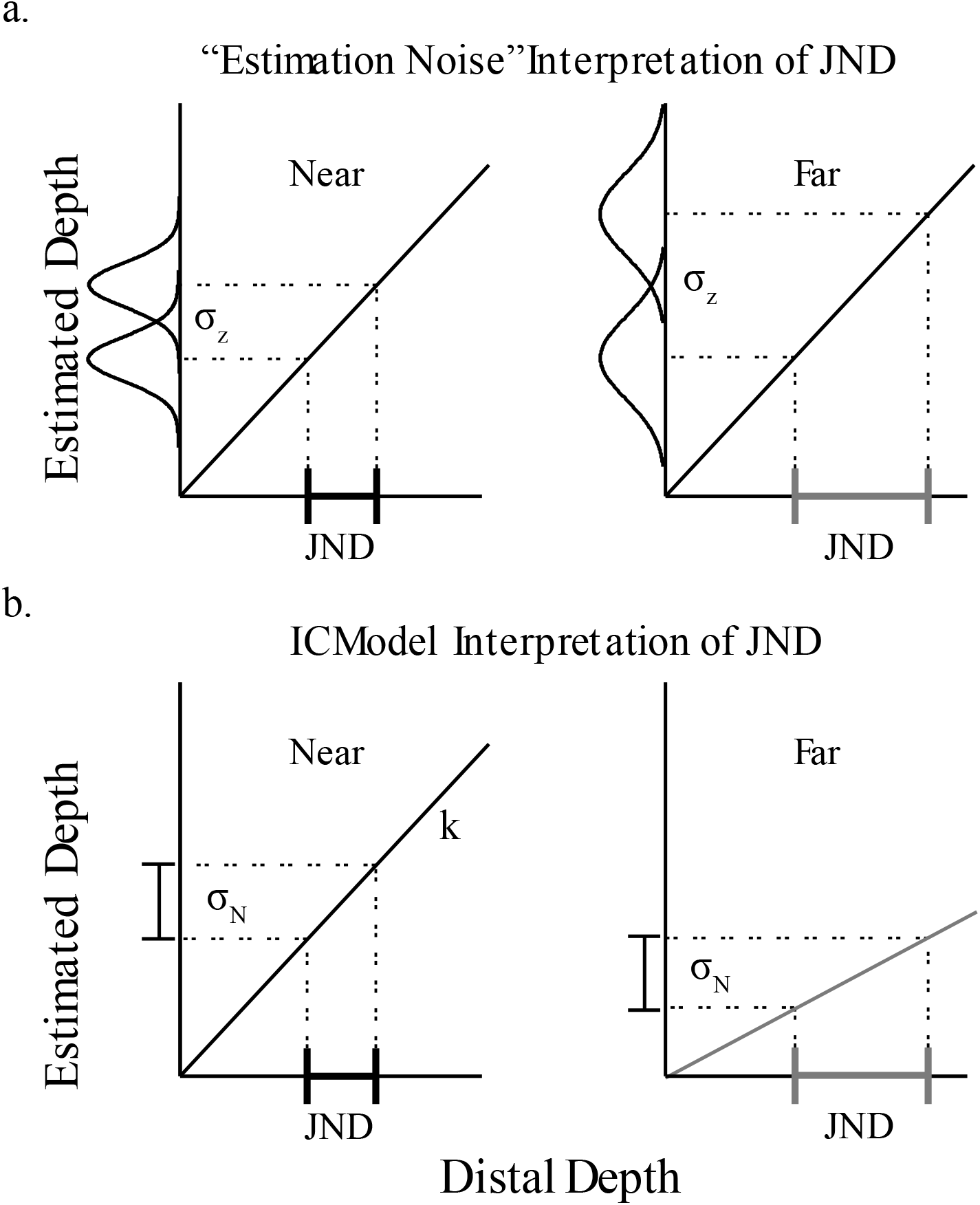
Two different interpretations of JND according to Bayesian theories (**a**) and the IC theory (**b**). **a**. Bayesian theories assume that variability of depth judgments are due to uncertainty of 3D estimates. For instance, disparities at near distances (left) are more reliable than disparities at far distances (right). Therefore the distribution of depth estimates are narrower at near distances than at far distances. In the example, only a small change of 5mm in distal depth is necessary to overcome the perceptual noise at near distances. However, at far distances a change of 10mm is needed. Note that the function relating distal depth to estimated depth is veridical. **b**. The IC theory predicts nearly deterministic estimates. However, it also predicts that the main cause of variability of perceptual judgements is task related. For the IC theory the JND measures the depth difference needed to overcome the task related noise. When the cue-strength is large, as what happens for disparity fields at near distances, only a small distal depth difference is needed. When the cue-strength is small, as it is for disparity fields at far distances, a larger distal depth difference is needed.

In contrast, the IC theory assumes that the noise in the estimation process is negligible. In other words, perceived depth is approximately the same across repeated viewings of the same stimulus under the same viewing conditions. However, noise in the response distributions of a task (*task noise*; often neglected by MLE models of cue combination) may arise due to factors such as response execution and memory requirements. Importantly, this noise is independent of distal stimulus properties such as texture quality or viewing distance. This leads to a different interpretation of the JND: given a particular cue strength, the JND is the change in distal stimulus magnitude needed to produce a perceptual difference that is large enough to overcome the effects of task noise *σ*_*N*_. As shown in the hypothetical experiment of Figure 6b, the JND is larger at the far viewing distance because the cue strength becomes weaker (consistent with the fact that binocular disparities decrease with viewing distance). We see that the JND is inversely proportional to the cue strength 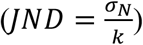. Recall that the Vector Sum model posits that adding cues to a stimulus increases the combined-cue strength according to the magnitude of the vector sum. Since the JND is inversely proportional to cue strength, the Vector Sum model therefore predicts that the JND shrinks with additional cues, similar to the MLE model. Specifically, the single-cue and combined-cue JNDs for stim uli defined by texture and/or disparity cues are given by 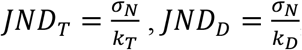, and 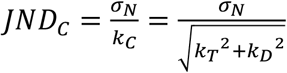. Appendix 3 shows how, from these equations, we can predict the combined-cue JND directly from the single-cue JNDs as follows: 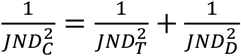 Notice that this equation is formally identical to Equation 2 of the MLE model, where JNDs are assumed to measure the estimation noise (i.e., *JND*_*i*_ = *σ*_*i*_). However, the Vector Sum model predicts that this relationship will hold at the same *perceived depth* (in order to equate task-related task noise, as the decision process operates on perceived depth), whereas the MLE model predicts it will hold at the same *simulated depth* (in order to equate estimation noise). Thus, the predictions of the two models for a given dataset may slightly differ, as we will show.

The goal of Experiment 2 was to demonstrate that the Vector Sum model correctly predicts the relationship between single-cue and combined-cue JNDs for the same stimuli presented in Experiment 1. Additionally, we aimed to show that this relationship is consistent with the independently measured cue strengths obtained in Experiment 1. These findings demonstrate that the IC theory’s interpretation of the JND is highly consistent with empirical results of depth discrimination tasks.

### *Experiment 2:* Methods

#### Participants

Eight participants from Experiment 1 returned to complete Experiment 2, including two of the authors.

#### Stimuli

Stimuli were identical to those in Experiment 1. However, in this experiment participants did not provide a judgment of absolute perceived depth. Instead, they performed a 2IFC depth discrimination task. Note that to make quantitative predictions of JNDs from the Vector Sum model, the perceived depth must be matched across the single-cue and combined cue standards so that the task noise, which is dependent on perceived depth, is kept constant. Thus, we used data from Experiment 1 to infer, for each participant in each viewing condition, a set of three simulated depths for texture-only, disparity-only, and combined-cue stimuli that elicited the same perceived depth (Figure 7a, horizontal lines). These simulated depths served as the standard stimuli in the 2IFC tasks, around which the JND was measured. For each viewing distance, we defined a large standard and small standard. The perceived depth that defined the small standard was anchored by the cue that elicited the greatest response at a distal depth of 2.5 mm. For the representative participant depicted in Figure 6a, the small standard corresponded to a perceived depth of approximately 4.5 mm, as this was the greatest reported perceived depth at 2.5 mm of simulated depth. Similarly, the simulated depth values for the large standard stimuli were anchored by the smallest perceived depth for a simulated depth of 15 mm. The simulated depth values for the various standard stimuli were chosen by interpolation using second-order curvilinear fits (see Fig. 7a). Through this procedure we determined 12 standard stimuli (3 cues x 2 viewing distances x 2 perceived depths) to be used in a 2IFC depth-discrimination task.

**Figure 7:**
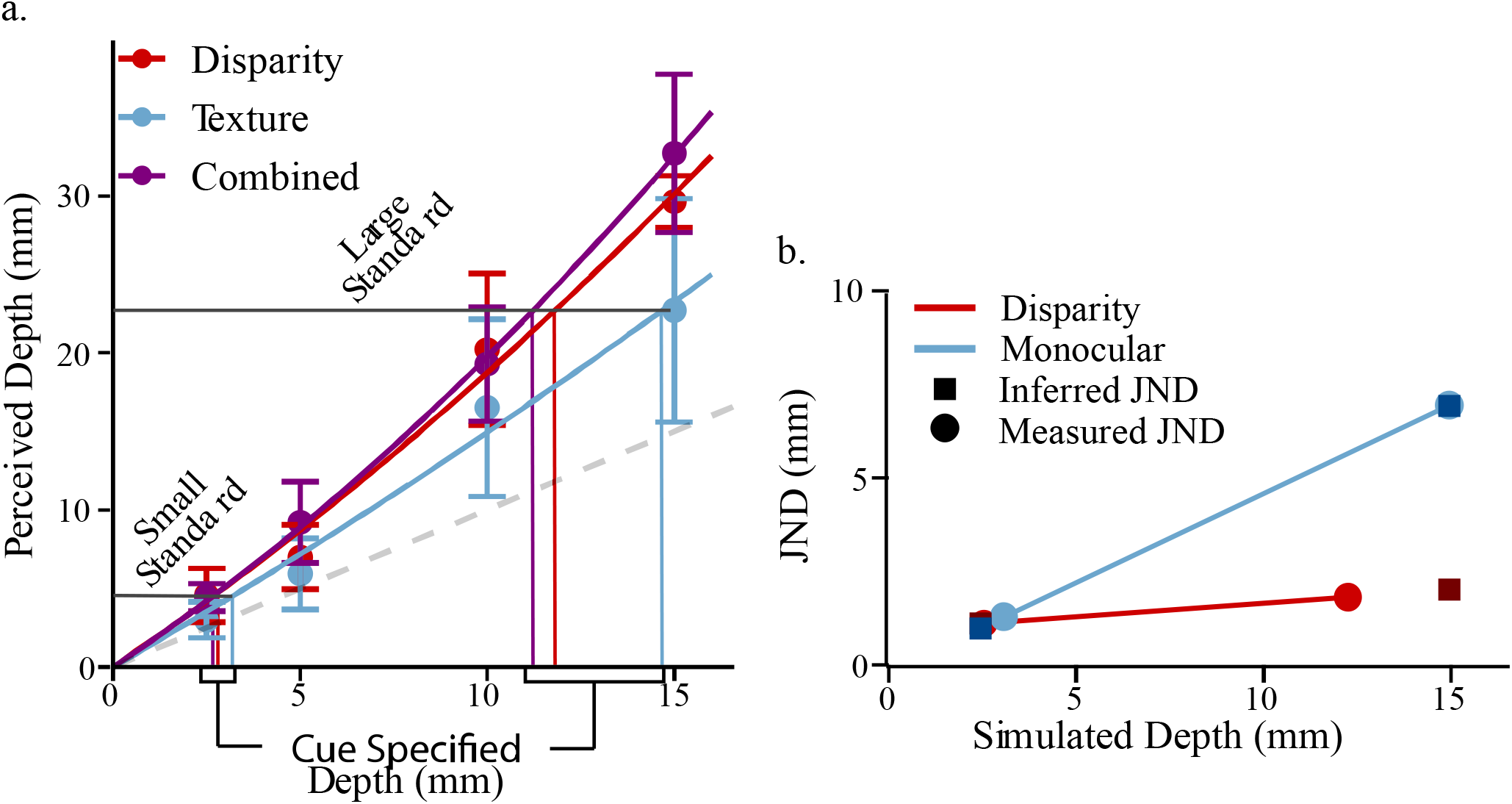
**a**. An example from a representative observer of how simulated depths for the fixed standards were chosen at 40 cm fixation distance. Two perceived depths were chosen, the largest and the smallest possible given the range of data. For a given perceived depth, each cue requires a different simulated depth to elicit that same perceived depth. These unique simulated depths were inferred for each cue through the intersection between curvilinear fits to the data (solid curved lines) and horizontal lines set at the preferred perceived depth. The vertical lines indicate these inferred values. **b**. MLE predictions need JNDs measured at the same simulated pedestal depth values. However, we measured the JNDs at slightly different pedestal values (solid circles). We therefore inferred the JNDs at the required pedestal values through interpolation or extrapolation (solid squares).

#### Procedure

Participants performed a 2IFC task in which the perceived depth of a standard stimulus with a fixed simulated depth was compared to that of a comparison stimulus whose simulated depth was varied through a staircase procedure. Four staircases were used in each condition (2-up-1-down, 1-up-2-down, 3-up-1-down, and 1-up-3-down) with 12 reversals each. On each trial, a fixation cross was displayed (700 ms), followed by the first stimulus (1000 ms), followed by a blank screen (1000 ms), then, again, the fixation cross (700 ms), and finally the second stimulus (1000 ms). Participants then reported with no time constraint which surface was perceived as having greater peak-to-trough depth through a keypress.

Response data were analyzed using a psychometric analysis package (Wichmann & Hill, 2001) in MATLAB. The data from each staircase procedure were fit with a cumulative Gaussian function. The point of subjective equality (PSE) was defined as the simulated depth at which participants responded with 50% accuracy. The JND was defined at the difference between the PSE and the simulated depth at which participants responded with 84% accuracy.

### Experiment 2: Results and Discussion

Figure 8 (colored bars) shows the average JND in each stimulus condition. On the horizontal axis, we indicate the average perceived depth corresponding to the two standard stimuli at each viewing distance. A repeated-measures ANOVA reported a significant main effect of cue (*F*(2, 14) = 25.42, *p* = 2.2e-5; Generalized *η*^2^ = 0.41). A critical prediction of both the MLE and Vector Sum model is that the combined-cue elicits a smaller JND than the single cue conditions. Bonferroni-corrected *t*-tests confirmed that the JND for the combined-cue stimuli (purple) was smaller than the JND for the disparity-only (red) (*t*(7) = -4.60, *p* = 0.005) and texture-only stimuli (blue) (*t*(7) = -7.93, *p* < 1.9e-4) conditions. Additionally, we found a significant main effect of perceived depth (*F*(1, 7) = 55.54, *p* = 1.4e-4; Generalized *η*^2^ = 0.38) with JNDs increasing for larger perceived depths. We suspect that this may be due to a form of Weber’s Law where the noise from the encoding and decoding of perceived depth to and from memory depends on the magnitude of perceived depth. We explore the implications of Weber’s Law further in the next section.

**Figure 8:**
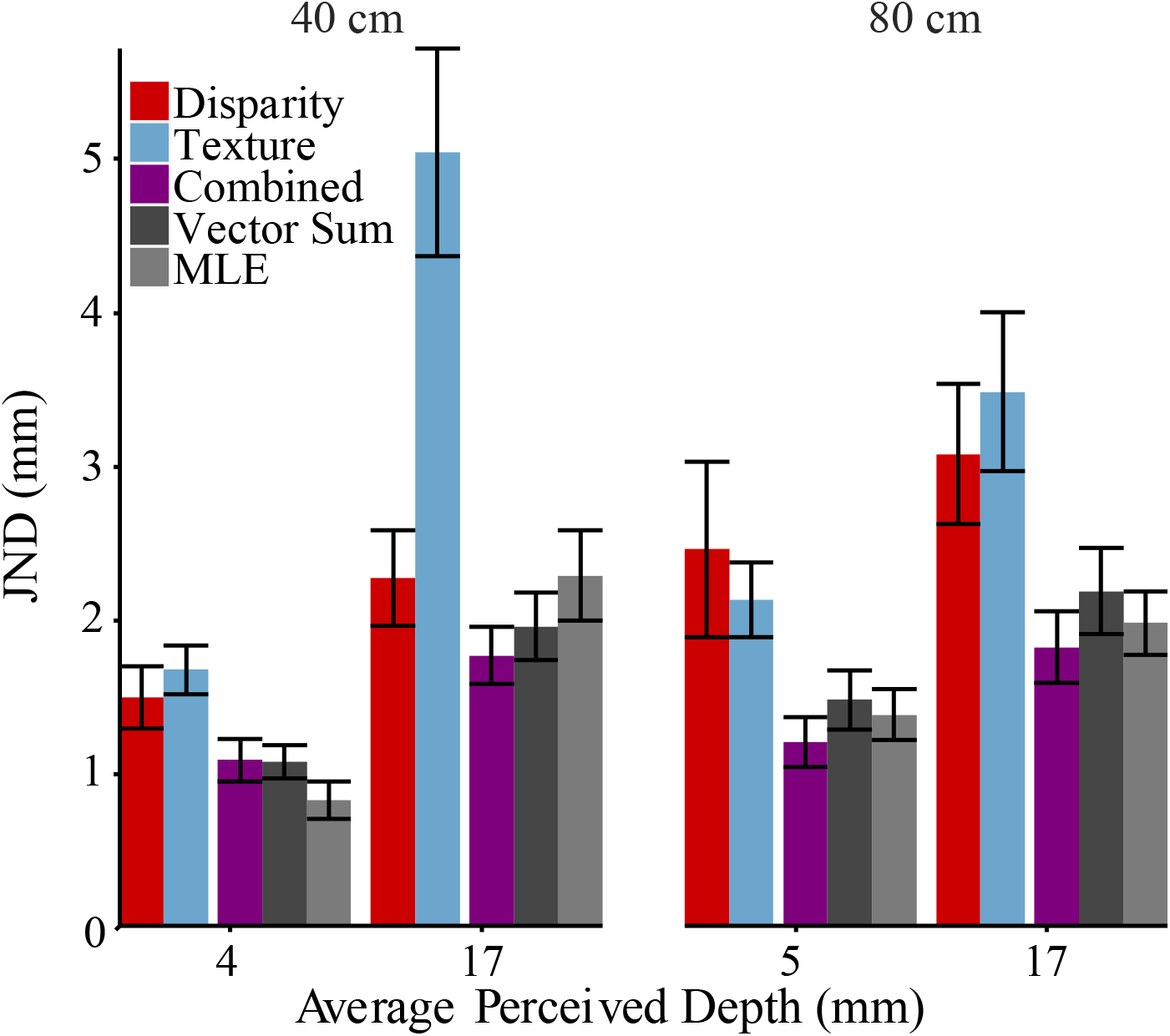
The Just-Noticeable Difference averaged across participants along with model predictions. The horizontal axis displays the average perceived depth of the standard. The perceived depth of the standard was chosen uniquely for each subject based on procedures stated in the previous section (See Figure 7a). The vertical axis represents the JND magnitude. Red, blue, and purple show JNDs measured for the disparity-only, texture-only, and the combined cue, respectively. Dark grey represents the Vector Sum model predictions while light grey represents MLE predictions. Error bars show standard error around the between-subject averages.

We also found significant interactions between perceived depth and viewing distance (*F*(1, 7) = 8.17, *p* = 0.024; Generalized *η*^2^ = 0.052), between cue type and perceived depth (*F*(2, 14) = 11.31, *p* = 0.0012; Generalized *η*^2^ = 0.20), and across all three factors of cue type, perceived depth and viewing distance (*F*(2, 14) = 5.54, *p* = 0.017; Generalized *η*^2^ = 0.074). These interactions, similarly, to Experiment 1, suggest a dependence of the cue strength on the cues and their viewing conditions. However, the key result is that the combined-cue JND is smaller than the single-cue JND in all conditions. Although this is often taken as evidence for the MLE model, here we show that it can also be predicted by the Vector Sum model.

The gray bars in Figure 8 show the predictions of the Vector Sum model (dark gray) and the MLE model (light gray) for the combined-cue JND. Recall that the Vector Sum model predictions are based on the single-cue JNDs for standard simulated depths that elicit the same *perceived depth* as the combined-cue stimulus. As mentioned above, this guarantees that the task noise was approximately matched across the three cue conditions. In contrast, the MLE model predictions are based on the single-cue JNDs for single-cue stimuli with the same *simulated depths* as the combined-cue stimulus. Although we did not measure the single-cue JNDs at fixed simulated depths, Figure 7b demonstrates how, for each participant, we linearly interpolated or extrapolated slightly from the measured JNDs (circles) to determine appropriate values for the MLE model (squares). Regardless, in Figure 8 we see that the predictions for the two models are very similar, as should be expected, with no significant difference in accuracy (*t*(7) = -0.39, *p* = 0.71).

#### Relationship Between JND and cue strength

The IC theory introduces the idea that the JND is not primarily a measure of estimation noise (which is assumed to be negligible), but rather than the noise that emerges from task-related demands involved in comparing two perceived depths across a time interval (*e*.*g*., temporal decay in memory). In a 2IFC task, the JND is determined by the cue strength of the varying comparison stimulus. This is because the cue strength determines how much change in the simulated depth of the comparison is necessary to produce a perceived depth difference large enough to overcome the task noise, *σ*_*N*_ (Fig. 6b). Thus, the JND depends on task noise and sensitivity to changes in distal depth. Furthermore, we expect that the JND is susceptible to Weber’s law, where increases in perceived depth will cause an increase in the standard deviation of the task noise. If we therefore assume that *σ*_*N*_ increases with the perceived depth 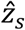of the standard stimulus through a Weber fraction *W*_*IC*_ then 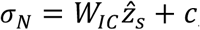, where *c* is aconstant reflecting a baseline noise. Since the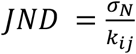, where *k*_*ij*_ is the cue strength of cue *i* (disparity, texture, and the combined-cue) for viewing condition j (40 cm and 80 cm fixation distance), we can obtain 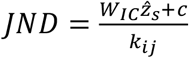. Because the perceived depth of the standard is 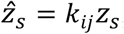where *z*_*s*_ is the distal depth of the standard stimulus, the JND can be modeled relative to the distal depth by Equation 6:

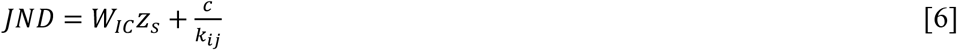

We expect that the JND depends (1) on the distal depth of the standard because of the Weber law, and, most critically, (2) on the cue-strength of the comparison *k*_*ij*_. We set, for each participant, the cue strength *k*_*ij*_ to the individual slopes from linear fits mapping the simulated depths observed in Experiment 1 to the perceived depths. To infer the Weber fraction and the noise coefficient, we fit Equation 6 to the estimated JNDs of each participant. We found both the Weber fraction (*M* = 0.13 mm, *SE* = 0.031 mm) and the noise coefficient (*M* = 1.66 mm, *SE* = 0.36 mm) to be significantly greater than 0 (*t*(7) = 4.11, *p* = 0.0045 and *t*(7) = 4.57, *p* = 0.0026 respectively). Critical here is that the JND measured in Experiment 2 depends on the cue strength observed in experiment 1 (Fig. 9a). Using Equation 6, we can discount for each participant from the observed JND the contribution of the Weber law and the constant reflecting the baseline noise so to produce a noise-corrected 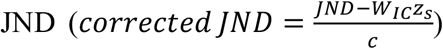. Figure 9b plots the relationship between the cue-strength and corrected JND averaged across participants. Horizontal error bars indicate the variability of the cue-strength across participants and vertical error bars the variability of the corrected JND across participants. Once the Weber fraction and the baseline noise constant are factored out, the JND is shown to be almost entirely dependent on the cue-strength 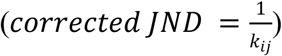 and independent of the cue type as predicted by the MLE model. For instance, the JND of the disparity stimulus at the close viewing distance (Fig. 9b, red circles) is smaller than the JND at the larger viewing distance (Fig. 9b, red triangles) because the strength of disparity at the smaller viewing distances is larger than the strength of disparity at the larger viewing distance.

**Figure 9:**
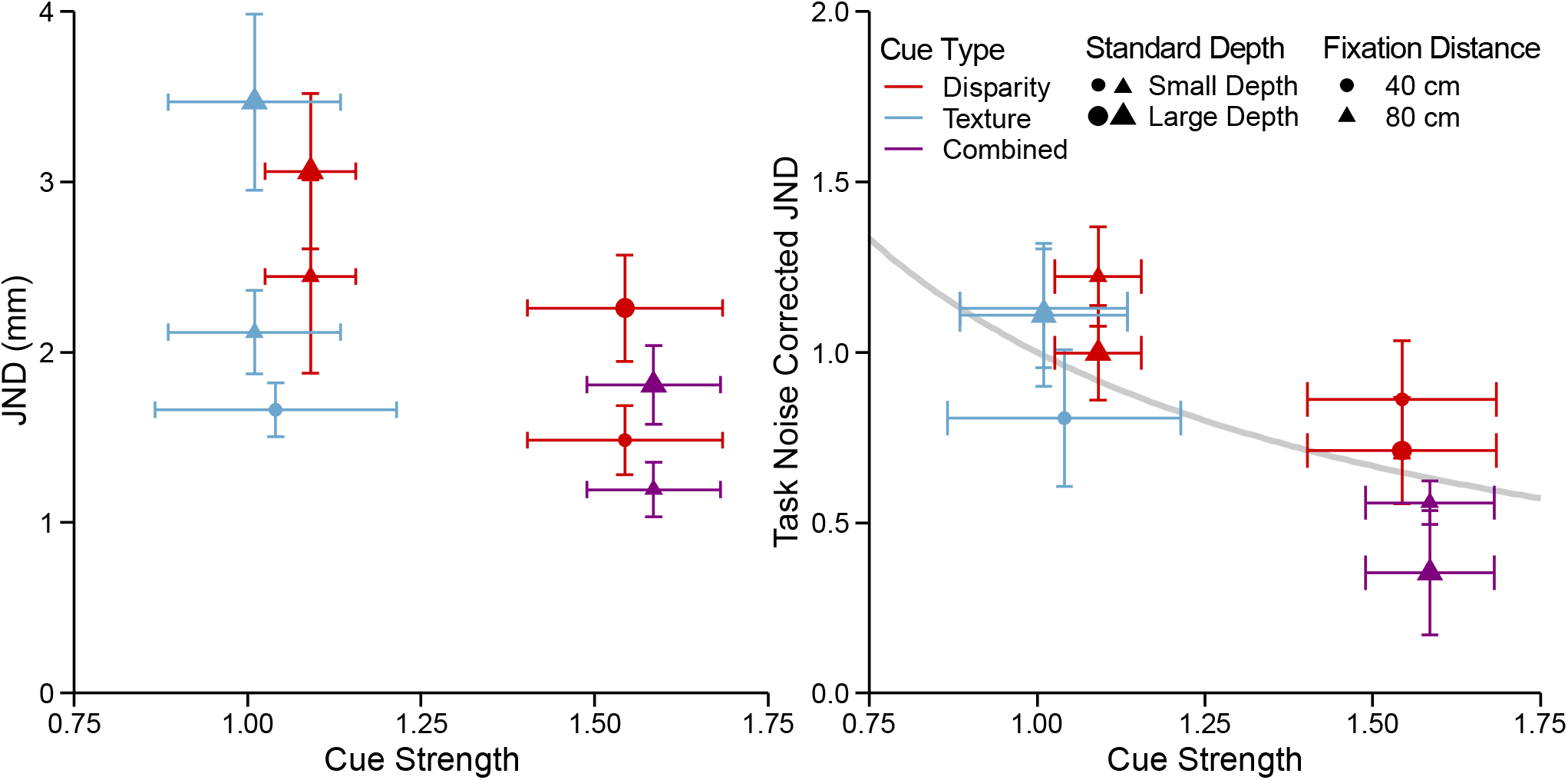
**a**. The JND against the cue strength cue strength extracted from Experiment 1 averaged across participants with *SE* bars. There is an inverse relationship between cue strength and the JND. **b**. Task noise corrected JNDs plotted against the cue-strength. The Vector Sum model predicts that there should be a hyperbolic relationship between the JND and the cue strength, which is plotted by the grey curve.

It should be noted that the condition for the large texture-only standard (average simulated depth of 17 mm) at 40 cm fixation distance was removed from this analysis for two reasons. First, we noticed that the JND in this condition is much larger than in the other conditions and, therefore, it constitutes an outlier (Fig. 8). Second, we also noticed that the function relating perceived depth to simulated depth for this condition is non-linear and seems to plateau at the largest simulated depth (Fig3, left panel, blue line). Because of this, the strength of the texture cue for larger depth values is smaller than the strength in correspondence to smaller depth values and, therefore, the JND at larger depth values is larger than the JND at smaller depth values.

In summary, these results indicate that the JND in a 2IFC t ask can be almost entirely explained by the cue-strength and not by the noise of depth estimates. This finding aligns with the predictions of the IC theory that postulates a deterministic mapping of depth modules between distal 3D properties and the module outputs.

## Experiment 3

The main aim of this experiment was to test a possible alternative interpretation of the results of Experiment 1, which are in agreement with the predictions of the Vector Sum model. The MLE model could accommodate the finding that combined-cue stimuli are perceived as deeper than single-cue stimuli once the role of “*cues-to-flatness”* is considered. Proponents of the MLE theory argue that when stimuli are rendered on flat displays, experimenters typically fail to eliminate all uncontrolled depth cues. As a result, residual depth information (*e*.*g*., the absence of a blur gradient) may specify the flat surface of the screen (Watt et al., 2005). If cues-to-flatness influence depth judgments, then single-cue conditions are inadvertently testing the combination of the single cue and the flatness cues. In this case, the MLE model predicts that the combined-cue stimulus may be perceived as deeper than the single-cue stimuli. Briefly, this is because the perceived depths of the single-cue stimuli are influenced more by the flatness cues than the combined-cue stimuli, due to differences in single-cue versus combined-cue reliabilities. On the other hand, the Vector Sum model directly predicts this well-known bias without postulating the influence of flatness cues. In fact, according to the Vector Sum model, flatness cues should have no influence on perceived depth because they specify zero depth and thus do not contribute to the vector sum.

In this experiment, we compared the two models predictions by testing whether intentionally adding flatness cues would reduce the perceived depth of a stimulus. In Experiment 3A, we compared perceived depth under monocular versus binocular viewing of the texture-only stimulus from Experiment 1. Binocular viewing of a texture-only stimulus with zero disparities provides a reliable flatness cue, akin to viewing a picture on a printed page, whereas monocular viewing of the same stimulus provides no such cue from disparities. Under the Vector Sum model, monocular and binocular viewing of a stimulus are equivalent, as they both have the effect of nullifying the disparity term in the Vector Sum equation (by setting either *k*_*D*_ = 0 or *z*_*D*_ = 0, respectively. Under the MLE model perceived depth should be greatly reduced under binocular viewing compared to monocular viewing, as disparities are posited to be highly reliable at near viewing distances, such that the disparity weight may exceed the texture weight. In Experiment 3B, we presented stimuli with the opposite relationship: binocular disparities provided non-zero depth information, but they were paired either with a textural flatness cue from a well-defined pattern specifying a fronto-parallel surface, or with an uninformative random-dot pattern often used to eliminate pictorial information from disparity-only stimuli. Here, the predictions are similar. The Vector Sum model predicts no difference in perceived depth, while the MLE model predicts a measurable difference.

Figure 10 illustrates the effects of cues-to-flatness for the MLE model for sinewave surfaces with either the uninformative random-dot pattern or the textural flatness cue. Figure 10c shows the predictions of the MLE model for the random-dot stimulus (Fig. 10a). As there is potentially some residual texture information from the random dots, this cue is represented as a zero-mean, large-variance distribution (blue). When combined with the reliable disparity cue (red), it has a negligible influence on the combined-cue estimate (purple). However, texture information is much more reliable for the polka-dot stimulus containing a textural cue-to-flatness (Figure 10b). Thus, in Figure 10d, the texture cue is represented as a zero-mean, small-variance distribution (blue). Consequently, when combined with the same reliable disparity cue (red), it will exert a larger influence on the combined-cue estimate (purple).

**Figure 10:**
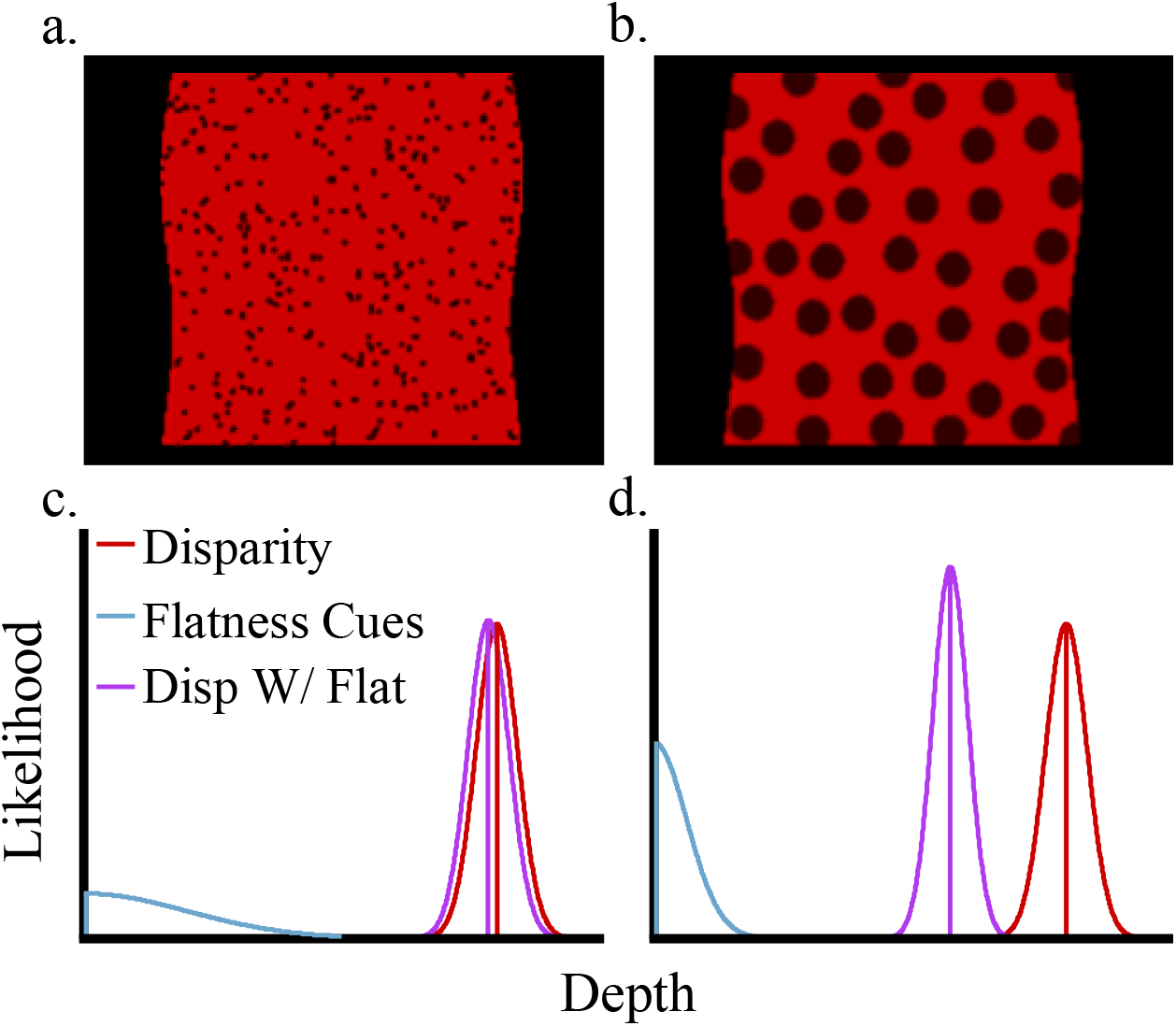
**a**,**c**. According to the MLE model, RDS displays (**a**) are expected to provide reliable depth estimates only from binocular disparity given the very low reliability of texture information (**b**). The red distribution represents the depth-from-disparity likelihood and the cyan distribution the likelihood of cues-to-flatness. The violet distribution shows the optimally combined distribution according to MLE. Note how the center of the distribution is only slightly pulled towards flatness. **b, d**. Unlike RDS displays, large circular polkadots (**b**) on the image plane reliably specify a flat frontoparallel surface. This flatness cue therefore produces a sharply peaked likelihood function centered at 0 depth (cyan). In this case the peak of the combined estimate is significantly pulled towards a flatter depth estimate (violet) (**d**). In contrast, the IC model predicts the same depth estimate in both conditions.

### Experiment 3: Methods

#### Participants

Seven observers participated in Experiment 3A, including two of the authors. Seven additional observers participated in Experiment 3B.

#### Apparatus

In Experiment 3A, the setup was the same as in Experiment 1, except that PLATO shutter glasses (Translucent Technologies Inc, Toronto, Ontario) were used to occlude the vision of the left eye during monocular viewing. Experiment 3B was conducted on a different system but using a similar setup (Alienware A51 with nVidia Quadro RTX 4000 GPU; Viewsonic G90fB CRT monitor, resolution 1280 × 1024, refresh rate 60 Hz; Volfoni Edge® RF controlled shutter glasses, Volfoni, Paris, FR).

#### Procedure

Stimuli and procedures were similar to Experiment 1 with a few exceptions.

In Experiment 3A, the corrugation in depth of the stimuli was specified by texture and shading cues (referred to as texture for simplicity; see Figure 3). However the same image was presented to the left and right eyes, producing zero disparities. Monocular and binocular viewing were randomly intermixed within the experiment, using the PLATO shutter glasses.

In Experiment 3B, participants judged the depth of a sinusoidal corrugation specified by disparity information in two conditions. The RDS (no-texture) condition was similar to the stereo-only condition of previous experiments, except the dots were painted black on a red background square subtending 8° of visual angle (along the diagonal) with an average of 292 visible dots. The dots subtended a visual angle of 0.05°. In the flat-texture condition we created a binocular stimulus that projected perfectly circular, 0.55° polka dots on the image screen by back-projecting the fronto-parallel texture onto the corrugated surface. Unlike Experiment 1, we also included the stimulus frame so that the only difference between conditions was the size and distribution of the texture elements.

##### Experiment 3A: Results

Figure 11 plots the average perceived depth as a function of simulated depth for monocular and binocular viewing at the two viewing distances. Repeated-measures ANOVA revealed a significant effect of simulated depth (*F*(1,6) = 49.94, p = 4.0e-4; Generalized *η*^2^ = 0.86). There were no other significant main effects or interactions. To evaluate the support for the Vector Sum model prediction of no difference between binocular and monocular viewing (*i*.*e*., the null hypothesis), we conducted a Bayes factor analysis. A Bayes factor of 0.21 indicated moderate evidence for a model including fixed effects of simulated depth and viewing distance and a random effect for participants, compared to a model including all three effects with an additional fixed effect of viewing condition. This supports the Vector Sum model prediction that the zero-disparity field specifying the flat picture plane does not influence perceived depth (see also Vishwanath and Hibbard, 2013). Overall, these findings seriously call into question the idea that the pattern of results observed in Experiment 1 (and in previous studies) is due to flatness cues. Moreover, the fact that depth perception is unaltered when viewing a pictorial stimulus with one or two eyes is successfully accounted for by the Vector Sum model.

**Figure 11:**
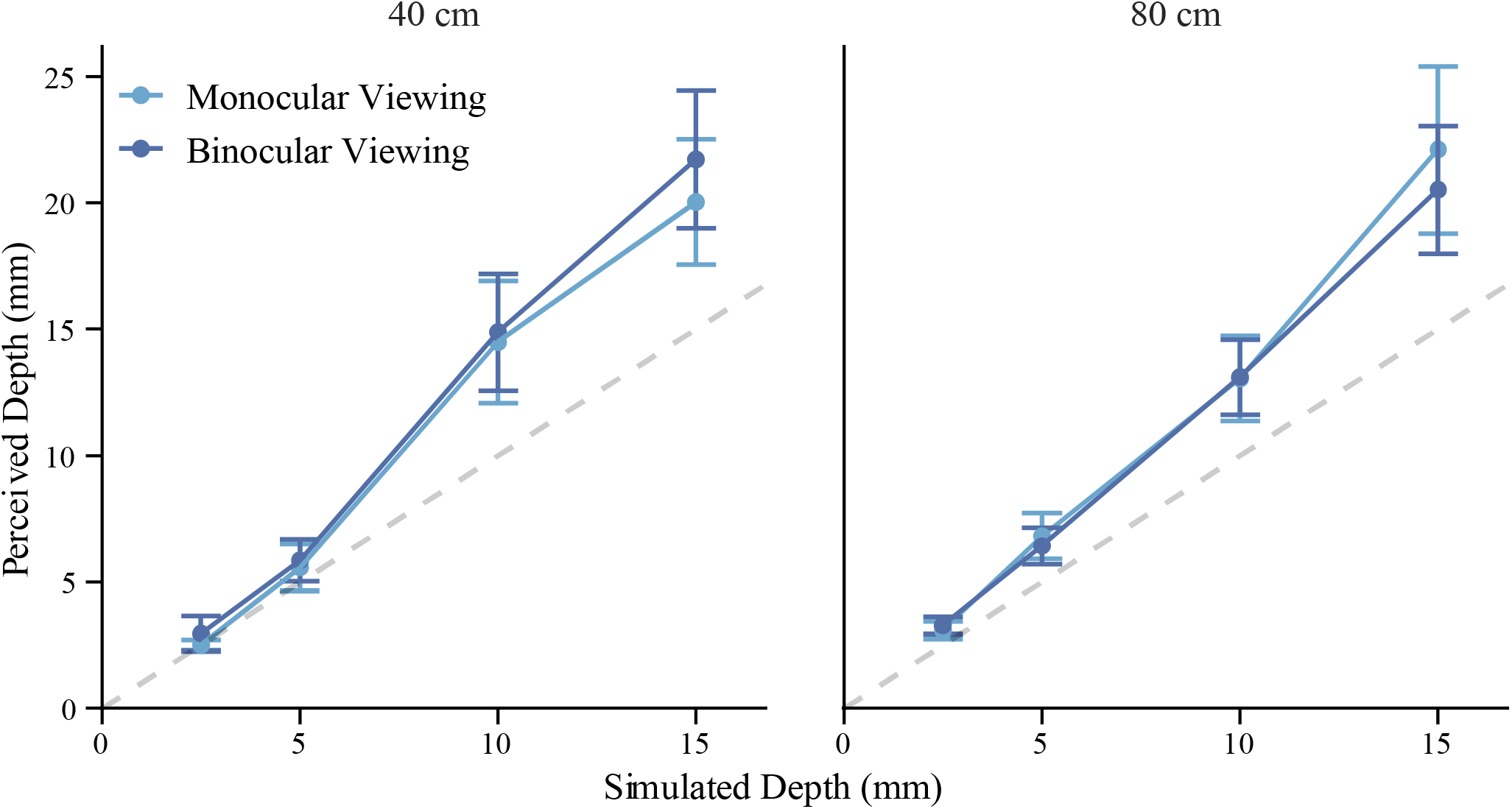
Averaged perceived depth as function of simulated depth in Experiment 3. 3D information is provided by texture and shading cues. Dark blue circles represent binocular view of the flat picture plane. Light blue squares represent monocular view. Error bars show the standard error around the average response.

##### Experiment 3B: Results

Figure 12 plots the perceived depth estimates in the flat-texture and random-dot conditions. Repeated-measures ANOVA revealed a significant main effect of simulated depth (*F*(1, 6) = 216.78, *p* = 6.2e-6; Generalized *η*^2^ = 0.92) and a significant interaction between simulated depth and fixation distance (*F*(1, 6) = 8.28, *p* = 0.028; Generalized *η*^2^ = 0.15). To evaluate the support for the Vector Sum model prediction of no difference between the flat-texture and random-dot stimuli, we again conducted a Bayes factor analysis. A Bayes factor of 0.42 indicated anecdotal evidence for a model including fixed effects of simulated depth and viewing distance and a random effect for participants, compared to a model including all three effects with an additional fixed effect of viewing condition. Together, the results of these experiments support the Vector Sum model prediction that there is no difference between setting the depth of a cue to zero or eliminating the cue altogether.

**Figure 12:**
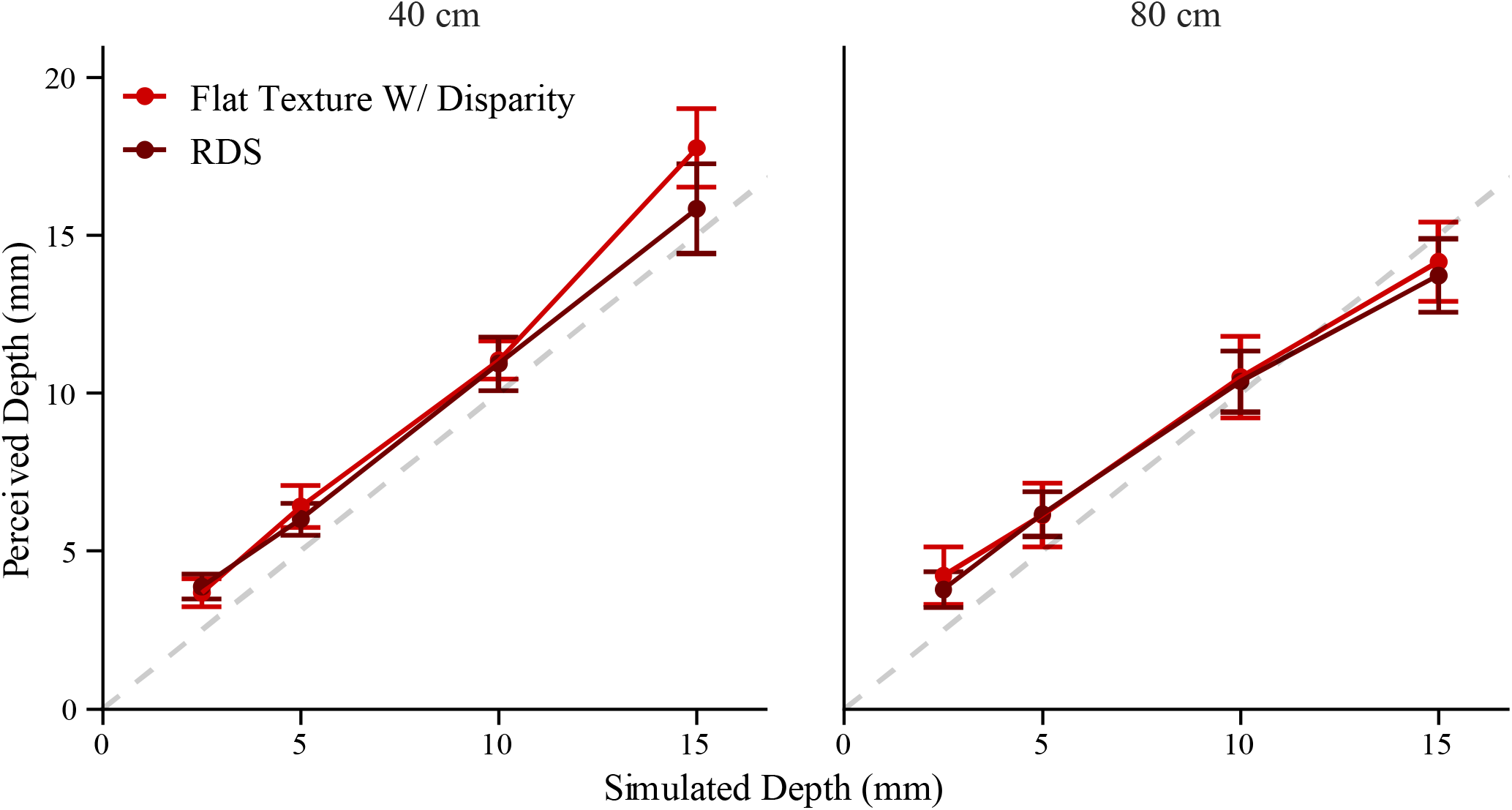
Averaged perceived depth as function of depth from binocular disparities in experiment 3b. Bright red indicates the RDS condition. Dark red indicates the viewing condition where polkadot texture specified a flat fronto-parallel surface. Error bars show the standard error around the average response.

## General Discussion

The results of three experiments challenge three fundamental assumptions of previous models of 3D cue integration. *Veridicality*: independent visual modules compute the veridical metric structure of 3D objects from retinal projections. *Probabilistic Inference*: the output of each module is a probability distribution of all possible 3D structures that may have generated a given retinal image. The width of these probability distributions is a measure of the perceptual estimation noise from each individual cue. In other words, each module has explicit access to information about the reliability of a given visual input. *Statistically Optimal Combination*: 3D cue estimates are optimally combined by computing the joint probability distribution from the independent probability distributions of each individual cue. The perceptual estimate corresponds to the 3D structure that maximizes this joint probability distribution. Moreover, since the joint probability distribution has a smaller variance than that of each individual cue the combined estimate is also more reliable. In the case of the linear MLE model, a simple heuristic can achieve *statistically optimal combination*: single-cue estimates are combined through a weighted average where the weights are inversely proportional to the variance of the noise of single cue estimates.

The *Veridicality* assumption is clearly contradicted by the results of the first experiment, where participants judged the amplitude of a surface with a sinusoidal depth profile. Following a classic cue-combination paradigm we studied these depth judgments with disparity-only, texture-only, and combined-cue stimuli. In most of these conditions the perceptual slopes relating simulated depth to perceived depth differ from unity and they significantly differ from each other. Moreover, the biases observed in single-cue conditions do not diminish when cues are combined.

The *Probabilistic Inference* assumption is challenged by the results of the second experiment where we show that JNDs measured in a depth discrimination task are inversely proportional to the slope of the transfer function independently measured in the first experiment. Since the perceptual slope is sufficient to predict depth-discrimination it presents a valid alternative interpretation of JNDs from the one postulated by the MLE models. Moreover, the IC theory’s explanation is more parsimonious since it does not assume mechanisms that have access to explicit measures of reliability of the visual input.

The *Statistically Optimal Combination* assumption is contradicted by the results of all three experiments. In the first experiment we found that the perceptual slope in the combined-cue condition is larger than the perceptual slope in the single-cue conditions, a result incompatible with the prediction of weighted cue combination of the linear MLE models. In the second experiment we show that the smaller JND in the combined-cue condition relative to the single-cue conditions can be explained by the larger perceptual slope. In the third experiment we show that adding a reliable cue-to-flatness to a 3D stimulus does not produce a significant reduction in depth magnitude. This finding contradicts the weighted cue combination rule of the MLE model, since adding to a depth cue a reliable cue-to-flatness should produce a weighted average that is biased towards flatness. These results should especially be expected when a flat disparity field is added to a texture specified 3D surface since at close distances disparity information is highly reliable. Instead, we observed no difference in perceived depth magnitudes when the picture of a texture stimulus was seen monocularly or binocularly. This finding also contradicts a possible MLE interpretation of the results of the first experiment. According to this interpretation, the larger slope of the combined-cue condition relative to the single-cue conditions may be attributed to the influence of spurious cues-to-flatness affecting stimuli rendered on flat CRT displays. The larger slope in the combined-cue condition is because these cues-to-flatness influence single-cue estimates to a greater extent than combined-cue estimates since the former are less reliable than the latter. If this explanation is correct and spurious cues-to-flatness such as the blurring gradient noticeably influence depth estimates, then we should expect an even larger effect when we introduce highly reliable cues-to-flatness such as a flat disparity field. But this is not what we found. In contrast to the observed discordance between the empirical data and the predictions of the MLE models, these findings can be accounted for by the Intrinsic Constraint theory of cue integration. These results therefore have significant theoretical implications since the IC theory rejects the fundamental hypotheses on which the MLE theory and the Bayesian approach in general stand.

### Linear mapping versus veridicality

The first important departure of the IC theory from previous theories is the rejection of metric accuracy as the normative goal of 3D processing. For the IC theory, mechanisms performing independent computations on the visual input derive 3D estimates that are *linearly* related to distal properties but are in general inaccurate. The slope of these linear functions, which we term *cue strength*, depends on the quality of the visual input. For instance, a regular pattern of texture elements on a distal surface such as polkadots will produce a stronger texture signal than sparse texture elements. Therefore, a depth-from-texture module will in the first case exhibit a steeper input-output transfer function than in the second case. Similarly, a disparity module will respond with a steeper transfer function to the depth of objects at closer distances than at further distances. The results of Experiment 1 show indeed that depth judgments are not veridical and depend on the viewing conditions. It can be observed that the perceptual slope in the disparity-only condition is shallower at a viewing distance of 80 cm than at 40 cm. At the smaller distance depth from disparity is overestimated and it is larger in magnitude than depth-from-texture. However, at the larger distance these estimates are almost the same.

### Deterministic versus probabilistic mapping

The second fundamental difference between the IC theory and MLE models is that the output of visual modules is *deterministic* and does not carry any information about the reliability of the input. Consider again a texture gradient projected by sparse surface texture elements. For the MLE account this is an *unreliable* image signal that produces a noisy output. In other words, each time similar (i.e. equally unreliable) stimuli are viewed the texture module will provide a different depth estimate. However, according to the veridicality assumption, the average estimate arising from multiple measurements will be unbiased. In contrast, the IC theory will derive similar depth estimates albeit much smaller than the distal depth magnitude. As explained above, what the MLE approach considers unreliable stimuli are considered as *weak* signals for the IC theory because a change in distal depth elicits a small change in the module output.

The deterministic nature of the mapping between distal and derived depth postulated by the IC theory requires an adequate re-interpretation of perceptual variability in depth estimation tasks. The most radical re-interpretation of variability measurements is with respect to the Just Noticeable Difference (JND) observed in depth-discrimination tasks. The MLE model considers the JND as a proxy measure of the standard deviation of the noise underlying perceptual estimates of depth. However, according to the IC theory, the noise influencing discriminability does not stem from variability of depth estimates, but, instead, from task processes. In the specific case of a 2IFC task, memory retention and retrieval of the stimulus presented in the first interval is subject to “smearing” (Rademaker et al., 2018), therefore affecting the following comparison with the stimulus presented in the second interval. To overcome this memory related noise the perceived depth magnitude of the two stimuli must differ by some minimum amount. Although this perceived depth difference necessary for a reliable discrimination is fixed, the *simulated* depth difference required to yield this perceived depth difference depends on the cue strength. Therefore, the JND, defined as the simulated depth difference necessary for a reliable discrimination, is inversely proportional to the cue strength. This novel interpretation of the JND is sufficient to predict the data of the second experiment since the observed JND is proportional to the inverse of the cue strength. Moreover, as we will discuss shortly, the Vector Sum rule of the IC theory and the alternative interpretation of discrimination thresholds yields the same prediction as the MLE model for the JND of combined-cue stimuli.

### Vector sum versus probabilistic inference

Within the IC framework independent depth modules have a deterministic input-output mapping. That is, the same type of visual input elicits the same output. However, this does not mean that the output of a 3D module is not subject to undesired fluctuations. The important distinction between the MLE theory and the IC theory resides in the nature of these fluctuations. For the MLE models the inferential process interpreting an unreliable visual input will produces large variations in the output estimates because even slight changes in the input will result in large perturbations of the associated likelihood function (Ernst & Banks, 2002; Held et al., 2012; Hillis et al., 2004; Knill 1998a,b; Knill, 2003; Knill & Saunders, 2003). It therefore makes intuitive sense that linear MLE models combine visual estimates with weights that are inversely related to the variance of the output noise. Note, however, that the weights must be estimated at each single instance and therefore visual modules must carry information about the reliability of a given visual input.

For the IC theory, fluctuations of a module output are caused by changes in the strength of the visual input. For instance, the same distal structure will yield 3D estimates of different magnitudes depending on the material composition of the object, the viewing distance, the illumination, and so on. It can be shown that the vector sum of the appropriately scaled module outputs minimizes the undesired influence of scene parameters while maximizing the sensitivity to distal depth changes (Appendix 1). This simple rule of cue combination yields specific predictions regarding both (1) the magnitude of depth judgments and (2) the discrimination thresholds of combined-cue stimuli. The first prediction is that the perceived magnitude of combined-cue stimuli is equal to the vector sum of the perceived magnitude of single-cue stimuli. Specifically, the cue strength (i.e. perceptual slope) of the combined-cue stimuli is the vector sum of the strengths of the single-cue stimuli. This prediction is confirmed by the results of the first experiment. The second prediction follows from the first. Since according to the IC theory, the JND is inversely proportional to the perceptual slope, it follows that the JND of the combined-cue stimuli is smaller than the JND of the single-cue stimuli (Appendix 3). Notably, the predicted reduction in magnitude of the JND for the combined-cue stimuli is identical to that of the MLE model. The algebraic equivalence of the Vector Sum and MLE prediction of the JND expected from cue combination validates the IC theory because it can account for many empirical findings that use depth discrimination to support the MLE predictions (Hillis et al., 2004; Knill & Saunders, 2003). Finally, the Vector Sum combination rule also predicts the results of the third experiment. When a cue to flatness is present in a display, its contribution to the vector sum is equivalent to that of an absent cue. For instance, when looking at a picture with only one eye, no disparity information is present whereas when looking at the same picture with two eyes the disparity field specifies zero depth. In both cases the contribution of the disparity term is nil.

## Conclusion

In this study we tested the predictions of a new theory of depth cue integration termed Intrinsic Constraint (IC) theory. This theory postulates the existence of independent modules relating perceived 3D properties to distal 3D properties through deterministic functions that are, in optimal conditions, linear. The slopes of these functions depend on scene parameters specific to the viewing conditions. In ideal viewing conditions depth modules are highly sensitive to distal changes in 3D properties, as for example when the material composition of an object determines a strong texture gradient. However, in viewing conditions where 3D information from a specific cue is weak, as for an object that only has very sparse texture elements on its surface, the response of the depth module will be shallow. The IC theory combines individual estimates through a vector sum that maximizes the response to changes in distal 3D properties while minimizing the module-output fluctuations due to varying scene parameters.

We tested this model in three experiments targeting different aspects of 3D shape judgments. First, we confirmed the prediction that increasing the number of cues specifying a 3D surface will increase the perceived depth of that surface, a hypothesis which we call the Vector Sum Model. This result has been recently found in other studies using grasping to test depth perception in both VR environments and with real objects (Campagnoli & Domini, 2019; 2022). Although Bayesian models can account for the phenomenon predicted by the Vector Sum model, the IC theory has the significant advantage of achieving the same predictions without the need for further ad-hoc assumptions such as cues-to-flatness or priors-to-flatness (Di Luca et al., 2010; Domini and Caudek, 2003, 2009, 2010, 2011, 2013; Domini et al., 2006; Domini et al., 2011). This advantage is not confined to the case of depth-cue integration, but it applies to other common visual experiences such as picture perception. In this case too, neither flatness cues nor a prior-to-flatness appear to be able to explain the empirical data. Second, we tested the ability of the IC theory to predict the JND of a multi-cue stimulus from the JNDs of single-cue stimuli. Notably, the IC theory makes formally identical predictions to those of Bayesian models, therefore accounting for a number of previous investigations that leverage JND data as the strongest source of evidence in support of Bayesian cue combination. However, the JND for the IC theory is determined by the slope of the response function and not by the noise of depth estimates.

In summary, the IC theory seems to be a better candidate for explaining 3D cue-integration experiments since (1) It can predict previous data in support of Bayesian models, (2) it predicts new results that are incompatible with previous models and (3) it is more parsimonious since it does not postulate veridical perception or needs estimates of cue-reliability that are necessary for the functioning of Bayesian models.

## Acknowledgements

The Authors would like to thank Dr. Dhanraj Vishwanath and Celine Aubuchon for their comments on a previous version of the manuscript and Kashif Ansari for his assistance in data collection. This work was supported by the National Science Foundation (Grant #1827550).

## Appendix

### Appendix 1: The Vector Sum equation maximizes the Signal-to-Noise-Ratio

For simplicity consider only two signals *s*_1_ = *λ*_1_*z* and *s*_2_ = *λ*_2_*z*, where *λ* _*i*_ are unknown multipliers depending on confounding variables and *z* is the magnitude of the 3D property. These signals are the visual systems encoding of the 3D information from independent cues (e.g. texture and disparity). We seek an estimate 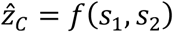 (1) Proportional to *z* and (2) Most sensitive to 3D information and least sensitive to random fluctuations *ε*_*i*_ of *λ*_*i*_. If *λ*_*i*0_ is the unperturbed value of *λ*_*i*_: *λ*_*i*_ = *λ*_*i*0_ + *ε*_*i*_ and *s*_*i*0_ = *λ*_*i*0_*z*. We assume small random perturbations due to changes in viewing conditions such that *ε*_*i*_ are Gaussian distributions with zero mean and standard deviations *σ*_*i*_. Taking the derivative of 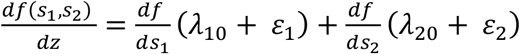, where 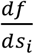 are calculated at *s*_*i0*_, we observe a signal term 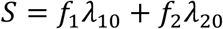 (where 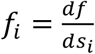) and a noise term *E* = *f*_1_ *ε*_1_ + *f*_2_ *ε*_2_ having standard deviation 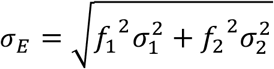. If we minimize the Noise to Signal Ratio 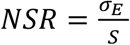 with respect to *f*_*i*_ (by solving for *f*_*i*_ the equation 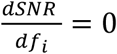) we find that the first derivatives of the function are 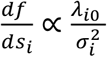. It can be shown that the derivatives 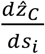 of the equation 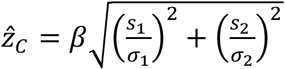 (calculated at *s*_*i0*_) meet this requirement. By substituting 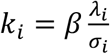 we obtain the Vector Sum equation 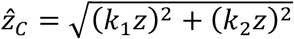 (easily generalizable to *n* signals).

### Appendix 2: The IC theory predicts the same linear combination rule as the Bayesian models in matching tasks

Hillis el al. (2004) predict the outcome of a task where the perceived slant of a non-conflict stimulus *S*_*NC*_ = *S*_*B*_ + *δ* is matched to that of a conflict stimulus *S*_*C*_, where *S*_*B*_ is an arbitrarily defined base slant and ?? is the change in slant needed for a perceptual match *E* (*Ŝ*_*C*_) = *E* (*Ŝ*_*NC*_). For the conflict stimulus the disparity slant *S*_*D*_ differs from a texture specified slant *S*_*T*_ by Δ: *S*_*T*_ = *S*_*B*_ and *S*_*D*_ = *S*_*B*_ + Δ. Optimal cue combination predicts that (*Ŝ*_*C*_) = *w*_*D*_(*S*_*B*_ + Δ) + (1 − *w*_*D*_)*S*_*B*_ = *S*_*B*_ + *w*_*D*_ Δ, where 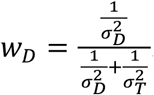. A matching (*E* (*Ŝ*_*C*_))= *E* (*Ŝ*_*NC*_)) is obtained when 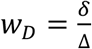 since *E* (*Ŝ*_*NC*_) = *S*_*B*_ + *δ*. By using JNDs as proxies for standard deviations the weight can be accurately predicted 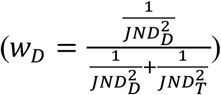. The IC theory makes identical predictions. For a small conflict Δ we can approximate the Vector Sum equation through Taylor expansion at the base slant *S*_*B*_: 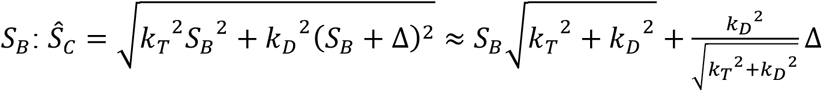. Since 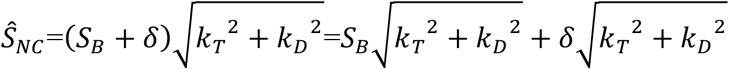, a match Ŝ_*NC*_ = Ŝ_*C*_ is obtained when 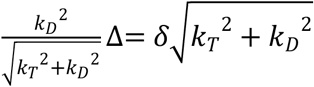, from which 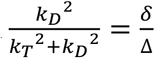. Note that since for the IC theory 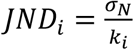 (See Introduction of Experiment 2) then 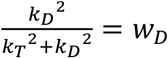, which matches Hillis et. al predictions.

### Appendix 3: The Vector Sum model predicts the same JND of combined stimuli as that predicted by linear MLE combination

The MLE model predicts that when two cues with independent Gaussian noise of standard deviation 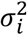are combined through a weighted average with weights inversely proportional to the variance of each cue then the combined (inverse) variance is 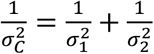. If JNDs are proxies for the standard deviations, then 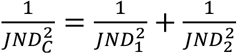. For the IC theory, JNDs depend on the task noise *σ* _*N*_ and the gain 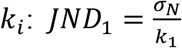and 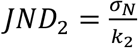. Since from the Vector Sum equation the gain of the combined stimulus is 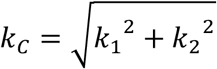, the JND of the combined stimulus is 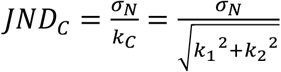. By substituting 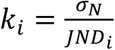 in this equation we obtain 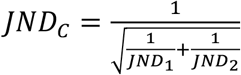, which is identical to the MLE prediction.

## Notes

**Conflicts of Interest** The authors declare no competing financial interests.

### Competing Interest Statement

The authors have declared no competing interest.

